# Mining Whole Genome Sequence data to efficiently attribute individuals to source populations

**DOI:** 10.1101/2020.02.03.932343

**Authors:** Francisco J. Pérez-Reche, Ovidiu Rotariu, Bruno S. Lopes, Ken J. Forbes, Norval J.C. Strachan

## Abstract

Whole genome sequence (WGS) data could transform our ability to attribute individuals to source populations. However, methods that effectively mine these data are yet to be developed. We present a minimal multilocus distance (MMD) method which rapidly deals with these large data sets as well as methods for optimally selecting loci. This was applied on WGS data to determine the source of human campylobacteriosis, the geographical origin of diverse biological species including humans and proteomic data to classify breast cancer tumours. The MMD method provides a highly accurate attribution which is computationally efficient for extended genotypes. These methods are generic, easy to implement for WGS and proteomic data and have wide application.

## Introduction

Attributing or assigning individuals to a source population is important within many disciplines including ecology, anthropology, infectious diseases and forensics^1,2^. For instance, assignment tests have been applied to identify the origin of individuals in ecosystems^3–7^, infectious diseases^8–19^, animals used for trade^5^ or the geographical origin of humans^20–22^ or plants^23^. A common strategy to attribute individuals to populations consists in comparing the genotype of the individual with the genetic profiles of defined source populations (e.g. the infectious disease example depicted in Fig. 1). The genotype usually comprises a set of genetic markers selected to highlight the differences between individuals (Fig. 1). For instance, highly variable genetic markers such as microsatellites^3–7^ or genes^8,11–19,24^ and more recently single nucleotide polymorphisms (SNPs)^25^ have been used for source attribution. The question is to decide which approach is most appropriate for the particular problem in terms of computation time and assignment accuracy.

**Figure 1.**
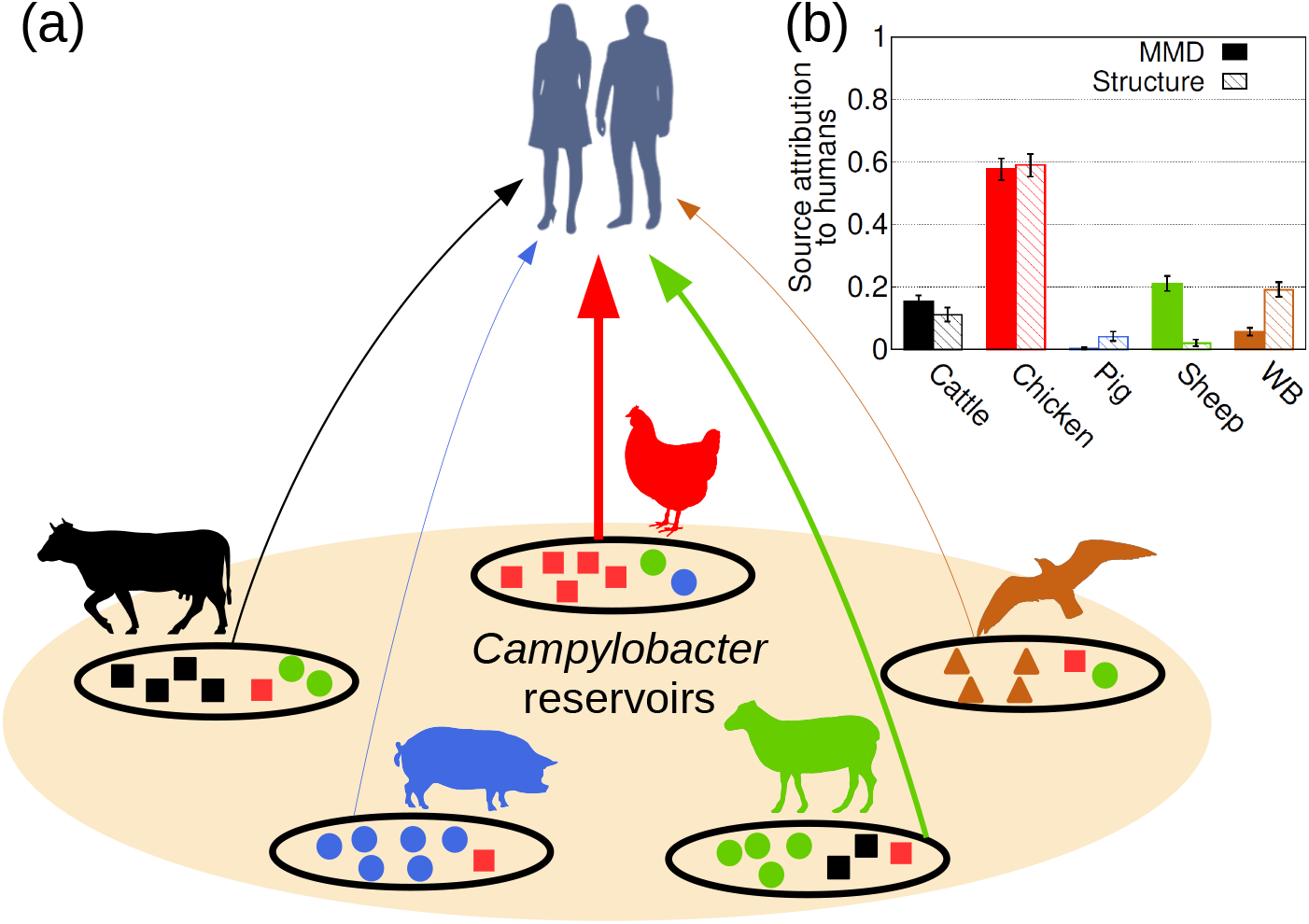
Source attribution. The general aim of source attribution (or assignment tests) is to determine the probability *p_u,s_* that an individual of unknown origin, u, originates from a certain source, *s*. Panel (a) provides the set of source populations considered in this study for *Campylobacter*: cattle, chicken, pigs, sheep and wild birds. The genetic profile of a source population *s* is represented by the genotypes of a set of *I_s_* individuals sampled from the source. Different symbols within sources schematically depict different genotypes. The genotype is determined from a set of genetic markers (loci) that depends on the typing method. Panel (b) provides the probability *p_s_* that any of the 500 human *Campylobacter* isolates are attributed to a source *s*. Results are shown for both MMD (solid bars) and STRUCTURE (hatched bars) methods, based on 25 937 cgSNP genotypes.

With the advent of next-generation sequencing technology, whole genome sequences are becoming available across all the 6 kingdoms of life ranging in size from for example viruses (kBases) to humans (3.2 GBases)^26–30^. In principle, this should enable discovery of large numbers of markers (e.g. SNPs) which have the potential to achieve unprecedented source attribution accuracy^31–36^. The challenge lies in efficiently mining large data sets for source attribution. Existing source attribution methods (e.g. STRUCTURE^37,38^ that has been widely applied in population genetics) operate on relatively short genotypes consisting of a few to tens or hundreds of loci. However, their computation time increases at least linearly with the number of loci and using extended genotypes (e.g. > 1000 loci) is impractical. This is a particularly important drawback in situations where rapid source attribution is crucial (e.g. for infectious diseases). To deal with extended genotypes, one either needs to develop efficient methods for source attribution and/or select sets of markers with high assignment power to keep the size of genotypes at a manageable level.

The limited effort to optimise source attribution algorithms to use extended genotypes contrasts with the effort made to address another important challenge in population structure, namely the use of extended genotypes to clustering individuals into groups. For instance, FRAPPE^39^, ADMIXTURE^40^, fastStructure^41^, fineStructure^42^, sNMF^43^, snapclust^44^, principal components analysis (PCA)^45^ or Discriminant analysis of principal components (DAPC)^46^ are well-known methods that can identify clusters using extended genotypes. In the language of machine learning^47^, these programs use unsupervised learning algorithms to infer clusters in the data without using any prior information about the characteristics of such clusters. Hence, such algorithms are not suitable for source attribution which requires supervised learning algorithms to classify individuals to a set of predefined sources whose characteristics are defined in terms of genotypes of known origin (i.e. in terms of a training set). ADMIXTURE was originally proposed as a method for unsupervised model-based estimation of ancestry of unrelated individuals^40^. This is the most widely used version of ADMIXTURE but, in fact, it was extended^48^ for supervised learning in such a way that it can use prior knowledge on the population of origin of some individuals to infer the ancestry of other individuals. The supervised learning version of ADMIXTURE, however, was not designed to estimate the probability that individuals were sampled from a certain source, i.e., it was not designed to attribute individuals to sources but rather to infer their ancestry. In spite of that, one would expect some relationship between ancestry and source of individuals and it makes sense to explore the capability of ADMIXTURE as an attribution method (with applicability restricted to datasets consisting of SNP genotypes). GLOBETROTTER, another package to infer the ancestry of individuals, also has potential as a method for source attribution with extended SNP datasets^49^.

Besides developing efficient methods for source attribution, selection of loci with high discriminatory power can also help deal with the computational challenge posed by extended genotypes. Several methods have been proposed to rank markers according to their importance for source attribution based on the intuitive idea that highly polymorphic markers should allow for higher genetic differentiation^50^. This can be achieved by measuring the importance of loci with diversity indices (e.g. expected heterozygosity, fixation index or informativeness^5,7,21,51,52^). Other approaches propose focusing on the joint performance of sets of loci rather than considering performance of loci individually^53–55^. One would expect these approaches to be more appropriate than diversity-based methods when dealing with correlated markers (i.e. when linkage disequilibrium is important^56^). However, they are computationally intensive and impractical to deal with extended genotypes and do not always improve on diversity-based methods^7^.

Here, we address two of the challenges posed by extended genotypes for source attribution. First, we propose a fast method for source attribution which can deal with genotypes comprising thousands of loci with minimal computational effort. Second, we propose the use of information theory^57^ for the optimal selection of markers from extended genotypes. We demonstrate this through several examples. The first is in the field of infectious diseases and involves *Campylobacter*, the largest cause of human bacterial gastroenteritis in the developed world^58,59^. Here we attribute human cases to source reservoirs (e.g. chicken, cattle, sheep etc.). The second is in the area of human evolution and involves attributing humans to 7 reference regions (e.g. Africa, Europe, etc.) or 53 populations (e.g. Bedouin, Maya, etc.)^22,60,61^. The third example studies attribution of the giant Californian sea cucumber (*Parastichopus californicus*) to north/south subregions within the northeastern Pacific coastal region^62^. In the fourth example, we assign breast cancer tumours to three different subtypes (ERPR, Her2 and TN)^63^. The first three examples use genomic data and the breast cancer example uses proteomic data. The performance of our method for source attribution is compared to the current state of the art method STRUCTURE^37,38^. For extended human genotypes which are too computationally intensive for STRUCTURE, a comparison is made with the supervised learning ADMIXTURE method^48^ by assuming that the probability of attribution to a source can be identified using the ancestry coefficient corresponding to such a source.

## Results

### Source attribution with the MMD method

We propose the Minimal Multilocus Distance (MMD) method to estimate the probability *p_u,s_* that an individual *u* is attributed to a population source *s* based on the similarity between the genotype of the individual to be attributed and genotypes from the sources. The similarity between pairs of genotypes is quantified by the Hamming distance which simply gives the number of loci at which the genotypes differ^64^. The smaller the distance between genotypes, the larger the probability that they originate from the same source (see Methods). To test the accuracy of the MMD method, we studied self-attribution, a cross-validation method^65,66^ which consists in removing individuals from the source population and estimating the probability that they are correctly attributed to their source based on the remainder^5,12,13,51^ (Fig. 2).

**Figure 2.**
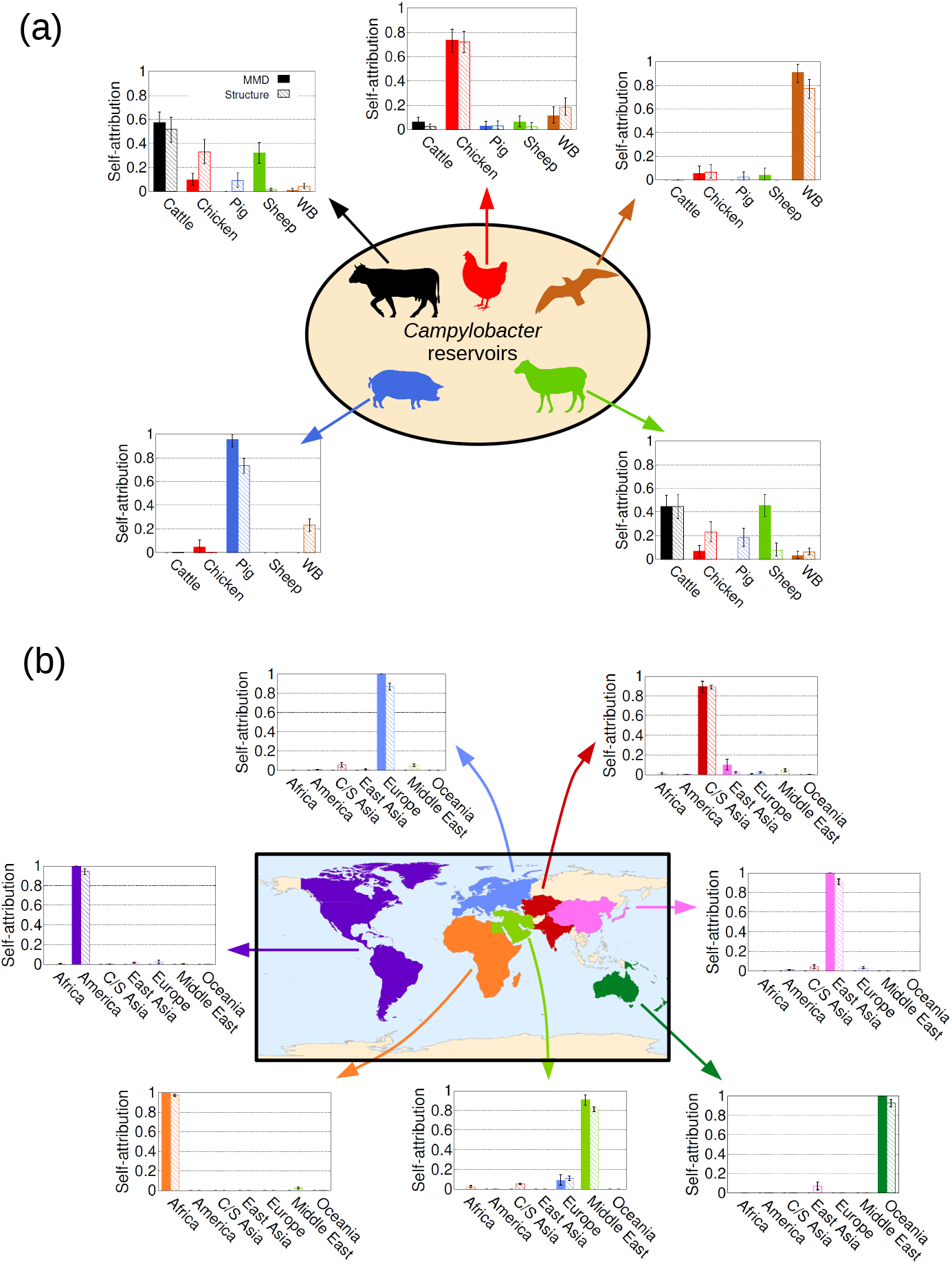
Self-attribution to test the accuracy of the source attribution methods. Self-attribution is a cross-validation strategy that involves removing individuals from the source populations and estimating the probability that they are correctly attributed to their source based on the remainder. Here 50% are removed. The bar charts provide the probability distribution *p_s_* for (a) *Campylobacter* (genotypes described by 25938 cgSNPs) and (b) humans (659 276 SNPs) comprising 5 and 7 source populations respectively. Bars in (a) show results obtained using the MMD method (solid bars) and STRUCTURE (hatched bars). Bars in (b) show results obtained using the MMD method (solid bars) and ADMIXTURE (hatched bars). Perfect self-attribution would result in 100% assignment to the appropriate source population. The total self-attribution accuracy when combining the results across all the source populations was, respectively, 73% and 56% for MMD and STRUCTURE in the *Campylobacter* population example. For the human population example, it was 97% and 71% for MMD and ADMIXTURE, respectively.

The source attribution results corresponding to a set of *I_u_* individuals (e.g. *I_u_* = 500 *Campylobacter* isolates from humans) are summarised by the probability distribution *p_s_* that any of the individuals is attributed to source *s* (see an example in Fig. 1(b) and more details in Methods). Self-attribution results are summarised by a similar probability distribution, assuming that the individuals that were removed from a population represent a set of *I_u_* individuals of unknown origin (see Fig. 2). In the following, we describe the results obtained for the *Campylobacter* and human examples. Self-attribution results for *P. californicus* genotypes and breast cancer proteomic data are described in Additional files 4 and 5, respectively.

#### Campylobacter

Self-attribution was carried out for isolates from food and animal sources by removing 50% of the isolates for blind attribution (Fig. 2(a)). Human clinical isolates are not considered for self-attribution since their source is unknown. The MMD method correctly assigned most isolates (> 70%) from pig, chicken and wild bird based on 25 937 core genome SNP (cgSNP) genotypes. Self-attribution of *Campylobacter* isolates from cattle and sheep is less precise (57% and 45%). Wrongly self-attributed cattle isolates are mostly assigned to sheep and chicken sources, whilst sheep isolates tend to be erroneously attributed to cattle and chicken sources. When combining the self-attribution results across all source populations, an overall attribution accuracy of 73% was obtained.

Source attribution was then carried out to predict the origin of the *Campylobacter* that resulted in human infection. As shown in Fig. 1, MMD estimated that most cases (61%) were associated with chicken whilst wild birds and pigs were relatively unimportant (< 8% for both sources). This is in line with a number of previous source attribution studies for human campylobacteriosis^11–19^.

#### Human

MMD self-attribution accuracy, based on removal of 50% of individuals genotyped at 659 276 SNPs^60^, was 100% accurate for all regions except for C/S Asia (90%) and Middle-East (91%). An overall self-attribution accuracy of 97% was obtained in this case (Fig. 2(b)).

Self-attribution based on 659 276 SNP genotypes was also studied at the level of the 53 populations available in the dataset^60^. In this case, an overall self-attribution accuracy of 73% was obtained. More explicitly, self-attribution accuracy was > 64% for 38 populations (see Fig. 3). Accuracy was poor for several populations from C/S Asia, E. Asia and Europe. For instance, individuals from the Uygur population (C/S Asia) were attributed to three populations in East Asia: Oroqen (40%), Hezhen (40%) and Japanese (20%). The attribution of individuals from some populations in East Asia (e.g. Mongola, Xibo, Cambodian, Han-NChina) was spread over other populations from East Asia. For the European region, individuals from French, Italian and Tuscan populations were often attributed to other geographically close populations. For instance, 17% of French individuals were correctly self-attributed, 39% were attributed to the Basque population, 17% to Orcadian, 13% to Sardinian, 10% to Italian and 4% to Tuscan.

**Figure 3.**
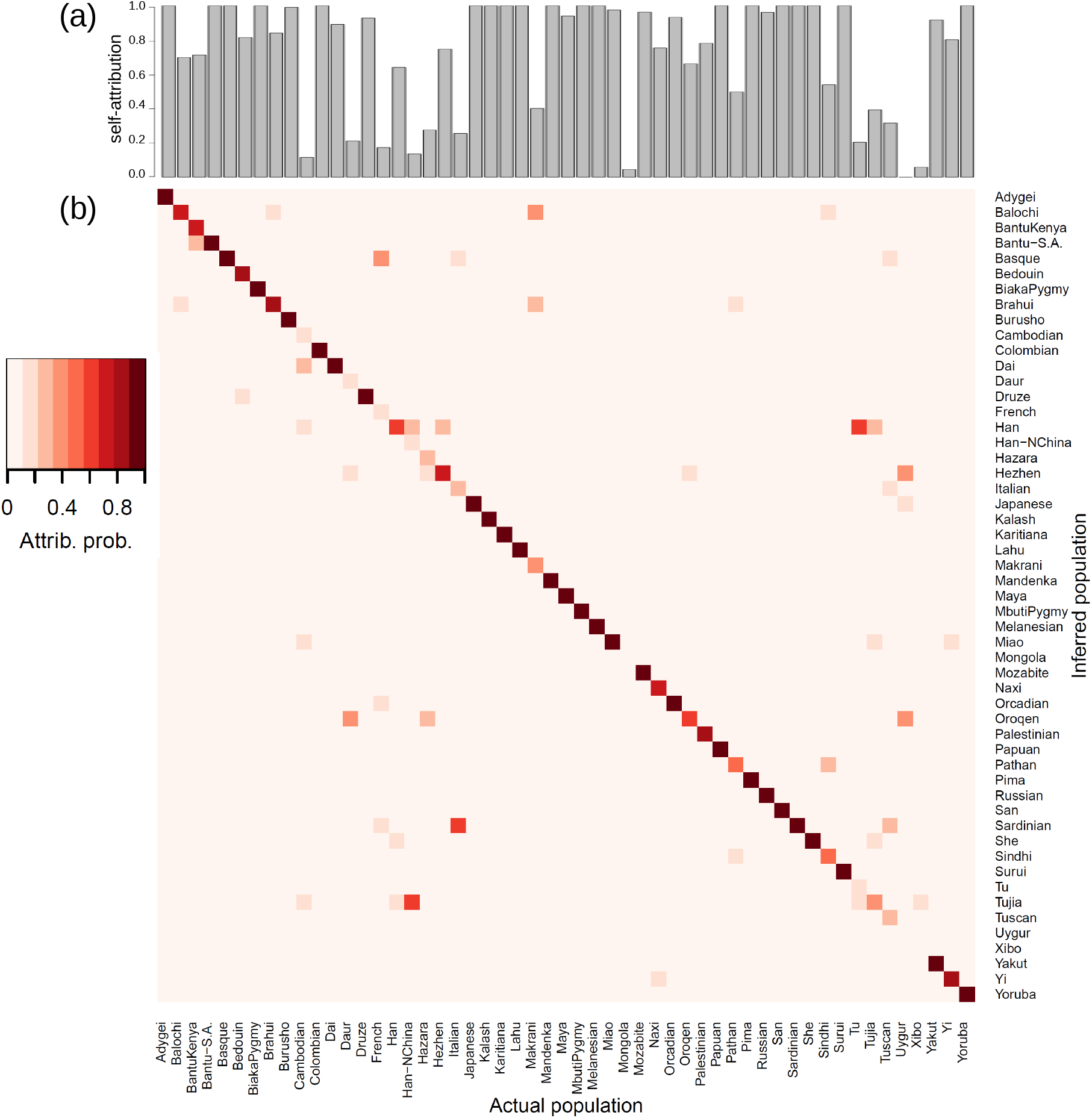
Self-attribution of humans to 53 sampling populations with MMD. Self-attribution for a given population was performed by randomly removing 50% of the individuals from that population and then attributing them to the populations characterized by the remaining individuals. (a) Probability of correct self-attribution for individuals selected from each population. (b) Attribution probability *p_u,s_* of removed individuals, *u*, to each of the populations, *s*. Darker colours correspond to higher probability, see the colour legend. The horizontal axis gives the population from which individuals were sampled and the vertical axis gives the inferred attribution probability to each population.

### Comparison with STRUCTURE and ADMIXTURE

The MMD method was compared with the current state of the art method STRUCTURE^37,38^ both in terms of attribution accuracy and computational speed. We used the *Campylobacter* and human genotypes for the comparison. Assuming that each source corresponds to a genetically-distinct population, STRUCTURE uses Bayesian inference to estimate the source attribution probability *p_u,s_* (see details on the implementation of STRUCTURE in Methods).

#### Campylobacter

Self-attribution tests for *Campylobacter* genotypes suggests that the probability of correct assignment calculated with the MMD method is higher than that obtained with STRUCTURE for all the reservoirs (overall 73% for MMD and 56% for STRUCTURE, see Fig. 2(a)). Both MMD and STRUCTURE have poorer self-attribution accuracy for cattle and sheep; the largest difference between MMD and STRUCTURE is observed for sheep isolates which are poorly attributed by STRUCTURE. In terms of source attribution of human *Campylobacter* isolates, both methods gave similar results with chicken being the most and pigs being the least important (Fig. 1(b)).

#### Human

A comparison of STRUCTURE and the MMD method based on extended 659 276 SNP human genotypes is not practical due to the long running time for STRUCTURE. In order to compare with STRUCTURE for humans, we considered smaller genotypes comprising 645 microsatellites^22^ and 2 810 SNPs^61^. For the microsatellite genotypes, MMD and STRUCTURE give similar overall self-attribution (87% compared with 84%, see Additional file 2: Fig. S1). Both MMD and STRUCTURE find it more difficult to differentiate between the European and Middle Eastern populations (Additional file 2: Fig. S1), due to a proportion of individuals in the European region being classified as Middle Eastern and vice-versa. When using the 2 810 SNP data set, STRUCTURE performed better with an overall attribution of 91% compared with 79% for MMD. The largest difference between MMD and STRUCTURE is observed for individuals from C/S Asia which are poorly attributed by the MMD method (Additional file 2: Fig. S2).

In order to compare the MMD method with existing methods for extended human genotypes comprising 659 276 SNPs, we run supervised analyses of ancestry using ADMIXTURE^48^ (see Methods for a more detailed description of ADMIXTURE implementation). The overall self-attribution accuracy achieved with ADMIXTURE is quite high (90%) but it is lower than for MMD (97%), see Fig. 2(b). In fact, self-attribution based on ADMIXTURE is less accurate than that obtained with MMD for all the regions. The largest differences between the self-attribution accuracy of the two methods were obtained for European individuals (100% with MMD and 87% with ADMIXTURE). This is mainly due to a significant contribution of C/S Asia and Middle East to the ancestry of Adygei individuals (see Additional file 2: Fig. S3). The self-attribution differences between the two methods are ≤ 10% for all regions except Europe. In particular, the smallest difference was observed for individuals from C/S Asia (90% with MMD and 89% with ADMIXTURE). In this case, MMD predicts a small probability of attribution to East Asia. ADMIXTURE predicts small probabilities of attribution to East Asia, Europe and Middle East (see Fig. 2(b) and Additional file 2: Fig. S3).

Our application of ADMIXTURE gives particularly low accuracies for attribution of individuals from Europe (49% correctly attributed), Africa (51%) and East Asia (67%). For several of the individuals selected from Europe, ADMIXTURE predicts a rather mixed ancestry from several regions other than Europe (see Additional file 2: Fig. S3). For instance, East Asian ancestry is inferred for some Italian, Tuscan and Adygei individuals. Middle Eastern ancestry is predicted for some Sardinian and Basque individuals. C/S Asian ancestry is predicted for some Basque and Russian individuals and African ancestry is predicted for Adygei, Russian and Sardinian individuals. For African individuals, ADMIXTURE predicts ancestry from Africa and, to a lower extent, from all the other regions except Oceania (Additional file 2: Fig. S3). In the case of individuals from East Asia, ADMIXTURE predicts significant ancestry from East Asia, Middle East and Europe.

#### Computational time

The computational time for MMD is much shorter than STRUCTURE. Fig. 4(c) shows a comparison of runtimes for self-attribution of *Campylobacter* isolates as a function of the number of SNP loci describing the genotypes. The MMD is between 100 and 10^5^ times faster than STRUCTURE for every run from 1 to ~ 2 × 10^4^ SNPs. Since the running time of MMD increases slowly with the number of loci compared to that of STRUCTURE, the efficiency of MMD improves relative to that of STRUCTURE for extended genotypes. For instance, STRUCTURE takes ~ 40h to assign a 25 937 cgSNP genotype whereas MMD completes the task in ~ 0.57 seconds (MMD implementation in R^67^, Processor: Intel^®^ Core^™^ i7-3770 3.40GHz).

**Figure 4.**
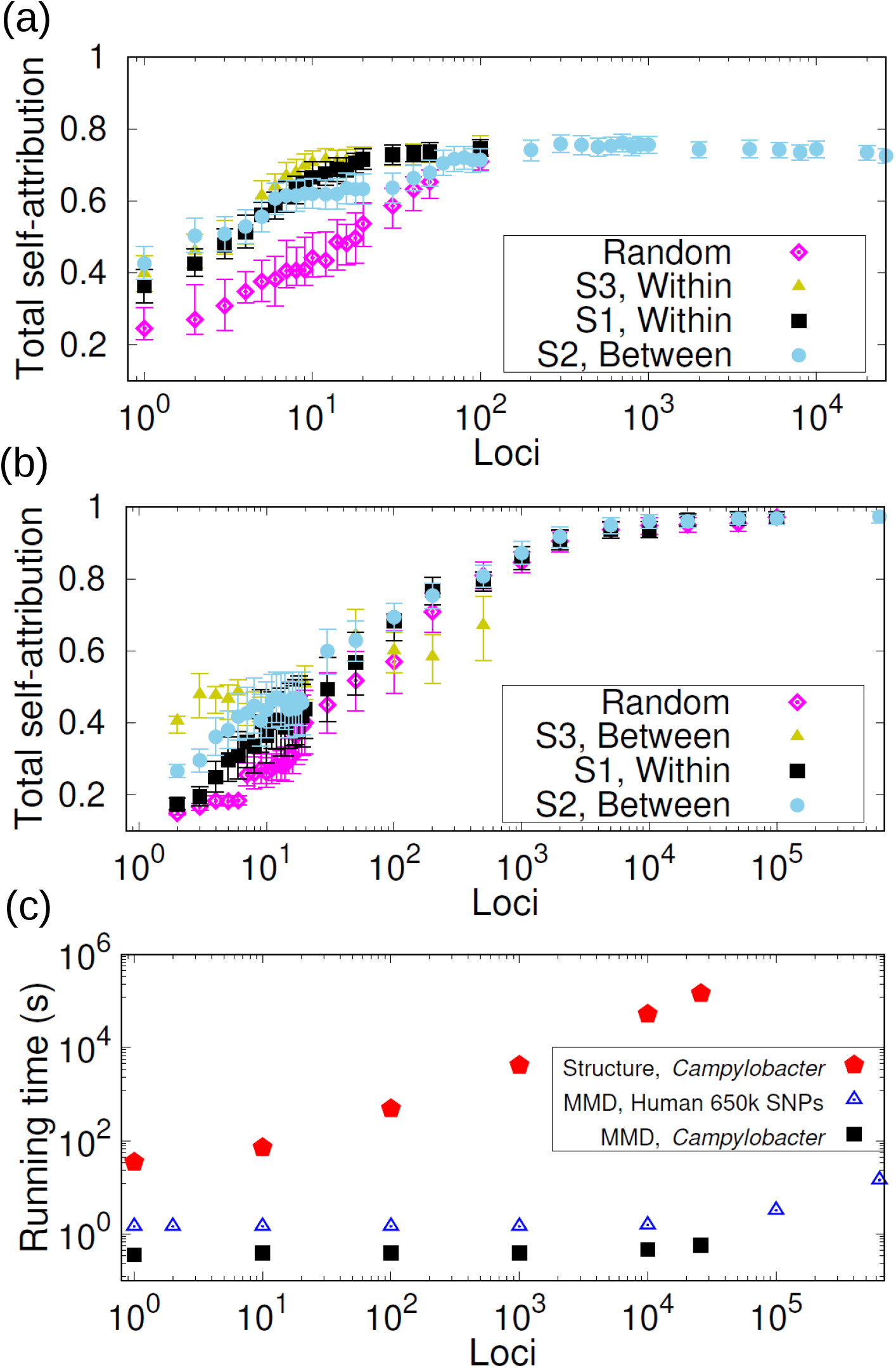
Selection of markers and computational times. (a) Total self-attribution probability p^sa^ that any *Campylobacter* isolate from food reservoirs is correctly attributed to its source. The probability is plotted as a function of the number of cgSNPs selected at random and with strategies S1 (loci ranked in decreasing within-source diversity), S2 (loci ranked in decreasing between-source diversity) and S3 (reordering the loci ranking corresponding to S1 to reduce loci redundancy). (b) Similar representation for the total self-attribution probability of human individuals based on 659 276 SNPs. Strategy S3 reorders the loci ranking corresponding to S2 to reduce redundancy. (c) Squares and pentagons show the computational time required for MMD and STRUCTURE, respectively, to assign a *Campylobacter* genotype with number of cgSNPs ranging between 1 and 25 937. Triangles show the time required for MMD to assign a human genotype with number of SNPs ranging between 1 to 659 276.

The MMD method is around twice as fast as ADMIXTURE when considering the 659 276 SNP dataset. More explicitly, MMD takes ~ 15 seconds to assign an individual whereas ADMIXTURE takes ~ 38 seconds to infer the ancestry of one individual (times based on an Intel^®^ Core^™^ i7-3770 3.40GHz processor for both algorithms).

For a given number of loci, the running time of MMD for attribution of a *Campylobacter* isolate is systematically smaller than the running time for attribution of a human individual from the 659 276 SNP dataset (see Fig. 4(c)). This is mostly due to the longer initial start-up time needed to deal with the human dataset which is significantly longer than that of *Campylobacter*. The running time remains essentially constant for runs with less than 10^5^ loci since the start-up time dominates over the MMD algorithm computational time which increases with the Hamming distance itself, not the number of loci (see Methods).

### Selecting informative markers for source attribution

In general, one would expect that the assignment power of a set of *n* markers will be subject to the following two conditions: (C1) Markers should allow us to capture the genetic differences between sources, i.e. the allele distribution of selected loci should significantly differ between sources. (C2) The *n* markers should contain complementary genetic information relevant to the source attribution process and ideally no redundant information. For example, the discriminatory power achieved with a set of markers will not increase significantly if a marker is added which brings redundant information compared to those that were already selected. Information theory offers a natural framework to account for allele diversity (relevant to C1) and loci redundancy (relevant to C2). Within this framework, allele diversity was quantified by the Shannon entropy and loci redundancy by the mutual information between pairs of loci (see Methods).

Inspired by these conditions, we propose *three strategies* to build sets of markers with high assignment power. Strategy S1 ranks loci in order of decreasing allele diversity *within* sources. Strategy S2 ranks loci in order of decreasing allele diversity *between* sources. Strategy S3 uses a greedy procedure^54^ that rearranges the loci obtained with S1 and S2 to reduce the redundancy. More explicitly, the list of selected loci in S3 is built by adding loci one by one making sure that the locus selected at the *n*-th step brings the smallest possible redundancy compared to the *n* − 1 previously selected loci. See the Methods section for an explicit definition of the redundancy *R^n^* of the *n*-th locus. Due to its greedy nature, strategy S3 is computationally more demanding than S1 and S2 and was applied to a limited number of loci to fine tune the selection of S1 or S2 (we only present the results of applying S3 to reorder the ranking given by the better performing strategy S1 or S2). The self-attribution performance of the three strategies as well as random selection of loci is illustrated in Fig. 4(a) for *Campylobacter*. As the number of selected SNPs increases, all of the targeted strategies (S1 to S3) saturated more quickly than random selection. The strategy that requires the fewest SNPs (approximately 10) to obtain optimal self-attribution is S1.

This is repeated for the human population examples. For the human 659 276 SNPs dataset, S2 does better than S1 for small numbers of loci but the difference becomes undistinguishable for *n* > 100 SNPs (see Fig. 4(b)). Strategy S3 brings some improvement over S2 for selections of *n* < 10 loci but does worse when selecting more loci. All tested strategies lead to saturation of the self-attribution accuracy when ~ 10^4^ SNPs are selected. For the microsatellite data set, strategy S2 also does better than S1 and strategy S3 does not improve on S2. Irrespective of the loci selection strategy, no sign of saturation of the total self-attribution is observed for the available loci (Additional file 2: Fig. S4). For the human 2 810 SNP genotypes, S2 does significantly better than S1 and S3 fractionally improves on S2 for selections of less than ~ 10 loci (Additional file 2: Fig. S5). However, the fraction of correctly self-attributed individuals increases slowly with the number of selected loci and using S3 does not represent a real advantage.

## Discussion

The source attribution problems studied here belong to a wider class of population structure challenges that also include classifying individuals in clusters of common features without assuming the population structure a priori. Significant effort has been made to optimise clustering algorithms to address the later challenge^39–46,48,68^. In contrast, optimisation of source attribution algorithms to use extended genotypes has received limited attention. The MMD method proposed here aims at filling this crucial gap.

The self-attribution accuracy of the MMD method is better than that of STRUCTURE for the *Campylobacter* example using SNPs (73% v 56%), approximately the same for the human origin from 7 geographical regions using microsatellites (87% v 84%) and slightly poorer than the human origin example using 2810 SNPs (79% v 91%). These results indicate its potential as an alternative to STRUCTURE. The MMD self-attribution accuracy of humans to 7 geographical regions increased to 97% when using 659276 SNPs. This is significantly better than the 71% self-attribution accuracy achieved by using the ancestry inferred by ADMIXTURE for the same dataset. A comparison with STRUCTURE was impractical for this dataset. The MMD method also gave a high self-attribution accuracy (73%) when using 659276 SNPs to assign humans to 53 populations.

Self-attribution of *Campylobacter* isolates from cattle and sheep reservoirs is poor compared to other reservoirs for both MMD and STRUCTURE methods. Similar trends have been reported in previous studies on *Campylobacter* self-attribution (see, e.g.^8^). This is likely due to the similarity of niche in cattle and sheep as both are ruminants. Also, geographical proximity offers frequent opportunities of transmission between the populations and this would explain the high genetic proximity between *Campylobacter* isolates from the cattle and sheep reservoirs (see Additional file 3: Table S1 where the allele-frequency divergence^69^ has been used as a measure of the genetic differentiation between sources).

Self-attribution of humans to 7 geographical regions based on microsatellites yielded lower accuracy for the European and Middle Eastern populations. This can be again explained in terms of the proximity of these regions, both geographically and genetically (see Additional file 3: Table S2 and^22^). The Central/South Asian population is also genetically close to the European and Middle Eastern populations but both the MMD and STRUCTURE methods provided a reasonably accurate self-attribution for individuals from C/S Asia. The human 2 810 SNP genotypes data set predicts a similar pattern for the allele-frequency divergence between populations (Additional file 3: Table S3); Europe, Middle East and C/S Asia are the genetically closest populations. Self-attribution of C/S Asian individuals based on the MMC method is, however, poorer for the 2 810 SNP data set than for microsatellites (compare Additional file 2: Figs. S1 and S2). Self-attribution accuracy increased when using the 659276 SNP dataset. In this case, self-attribution was very accurate (> 90%) for all 7 regions with only a 10% chance that individuals from C/S Asia and Middle East are wrongly attributed. This can again be ascribed to the relatively high genetic and geographical proximity between individuals from these regions (see Additional file 3: Table S4). Self-attribution of humans to 53 populations using the 659276 SNP dataset was highly accurate for most populations (overall accuracy of 73%). Populations that were poorly self-attributed are again genetically and geographically close to those populations to which they were wrongly attributed.

The self-attribution accuracy achieved for *P. californicus* with the MMD method is, within statistical error, comparable to that obtained in Ref.^62^ (see Additional file 4). The self-attribution accuracy of breast cancer tumours is relatively low (63% overall correct self-attribution, see Additional file 5). The fact that wrongly attributed samples are evenly attributed to the two wrong subtypes is likely due to the similarity between subtypes (see Additional file 3: Table S5). Our hypothesis is that the self-attribution accuracy could significantly improve by extending the dataset with more samples to describe the subtypes.

The fact that the MMD method uses the Hamming distance between genotypes contrasts with many other assignment methods that rely on allele frequencies^3,5,14,15,19,37,38,50,51,70–73^. This includes a range of methods that use frequency-based genetic distances that differ from the Hamming distance^51,74^. Using the Hamming distance makes the MMD method intrinsically faster than frequency-based methods. Indeed, the runtime complexity of frequency-based methods increases linearly with the number of loci in the multilocus genotypes. In contrast, the computational complexity of the MMD method increases with the Hamming distance (see Methods). Since the Hamming distance is typically smaller than the number of loci (see some examples in Additional file 2: Fig. S13), this represents a significant speed improvement.

Frequency-based assignment methods (including those using genetic distances) traditionally quantify the similarity between the individuals and sources in terms of a scalar quantity (e.g. a genetic distance or the value of a likelihood function, see Methods). In contrast, the MMD describes the similarity between individuals and sources in terms of the probability distribution of the distance (more explicitly, it uses the cumulative distribution function *F_u,s_*(*λ*) of the Hamming distance, as described in the Methods). Measures of similarity used in traditional methods could be regarded as summary statistics of the distribution function. For instance, for unlinked loci, the likelihood function used by some frequency-based methods^3,37,51,70,74,75^ corresponds to the probability that the Hamming distance between an individual and a source is zero (see Additional file 6). In general, the distance probability distribution gives a more complete description of the similarity between individuals and sources than specific characteristics of the distribution. The *Campylobacter* dataset is an interesting example in which using the whole distribution is convenient since it is often bimodal and a description in terms of a single statistical measure might not be appropriate (see Additional file 2: Fig. S13).

The MMD method assumes that the genetic profile of populations is defined by the genotypes of the individuals sampled from each source. In this respect, it is similar to some distance and frequency-based methods that determine the allele frequencies straight from the observed genotypes^3,15,50,51,72–74^. The frequency-based methods show a certain arbitrariness when an allele is present in the individual to be assigned but it was not observed in any of the sources. In order to make sure that the individual is assigned to a source, some methods set the frequency of the missing allele in the sources to a small value^76^ or to a value given by the inverse of a beta distribution^25^. In the MMD method, a missing allele will simply contribute one unit to the Hamming distance between the individual and all sources. The MMD method implicitly assumes that those alleles that are missing in all sources do not bring any relevant information for source attribution.

Strictly speaking, the allele probabilities of a population cannot be fully determined from the observed allele frequencies in a sample (i.e. the sample will typically not cover the whole population and observed allele frequencies only give an approximate representation of the genetic profile of the population). To circumvent this problem, several frequency-based assignment methods use Bayesian approaches to model the allele probability distributions of the populations^5,14,19,37,38,51,70,71^. It has been reported that source attribution based on Bayesian methods often outperforms plain frequency-based methods^51^. Extending the MMD method by using Bayesian methods to infer genotypes within sources is a possibility that could be explored in the future. However, since we are now immersed in the big data era, to take advantage of this it is likely that a better strategy to ensure high assignment accuracy can be achieved exploiting non-Bayesian techniques such as the MMD method.

The largest differences between MMD and STRUCTURE self-attribution results were observed for sheep *Campylobacter* isolates (STRUCTURE does poorly, see Fig. 2(a)) and humans from C/S Asia based on 2 810 SNPs (MMD does poorly, see Fig. S2). We hypothesise that these differences could be associated with two factors. On the one hand, STRUCTURE uses sophisticated methods to infer the allele probabilities of sources. In principle, such probabilities could give a more precise characterisation of sources than those used in the MMD method which is just based on observed genotypes. On the other hand, even small errors in the estimate of the allele probabilities for STRUCTURE lead to an attribution error that increases faster with the number of loci than that of the MMD method (according to our arguments in Additional file 6, this is expected for any method using a likelihood function to measure the similarity between individuals and sources^3,5,14,15,19,37,38,48,51,70–74^). Based on these considerations, one would expect a lower accuracy for the MMD method when using genotypes with a relatively small number of loci (e.g. for our 2 810 human SNPs example). In contrast, for extended genotypes, the error of the likelihood function used by STRUCTURE can become large and this may result in a poor attribution accuracy compared to that of the MMD method.

ADMIXTURE also uses a likelihood function to estimate the ancestry and allele frequencies and this might explain its lower self-attribution accuracy compared to MMD for the human 659 276 SNP dataset. In spite of that, ADMIXTURE gave a rather accurate self-attribution for this dataset and deviations from a perfect self-attribution can be explained in terms of geographic and genetic proximity between regions (e.g. the probabilities for attribution of European individuals to C/S Asia and Middle East).

In general, the performance of any method might depend on specific details of data sets (e.g. distribution of populations within the genotype space and the level of intermixing). Identifying the specificities of data sets that would favour one source attribution method over another in terms of accuracy can be achieved on a case by case basis employing training datasets as we have done here for self-attribution. However, this might require laborious analysis of genotypes to find specific features.

A central assumption of assignment methods is that the set of sampled sources includes the true population of the individual to be assigned. Accordingly, individuals are assigned to at least one source even if there is a big difference between the individual and all sources. The MMD method is not different in this respect. In order to assess the likelihood that the true population of origin of an individual has been sampled, one should use an exclusion test^51^. We applied the threshold exclusion method proposed in Ref.^5^ for STRUCTURE to the MMD attribution for human genotypes with 659276 SNPs and *Campylobacter* genotypes with 25 937 SNPs (see Additional file 7). The method only assigns an individual to a source if the attribution probability *p_u,s_* is above a threshold *T*. We found low exclusion rates for regions in the human dataset but exclusion was significant for the *Campylobacter* example even for self-attribution tests in which sources were definitely sampled. To understand the high exclusion rate for *Campylobacter* isolates, one should bear in mind that exclusion based on the threshold method does not necessarily imply that the source of the individual to be assigned has not been sampled. Instead, it might be a signature of a low genetic differentiation between sources. Consider, for instance, two genetically similar sources. The probability that an individual from one of the sources is attributed to any of the two sources will be around 1/2. Despite the fact that the source of the individual was definitely sampled, a threshold method will exclude both sources unless the threshold is very low (i.e. *T* < 1/2). When sources are not completely different to each other, it makes sense to assign individuals to several sources with certain probability rather than excluding sources with low assignment probability. For instance, the probabilistic assignment to several sources done by the MMD method should be the best way to capture the uncertainty in inference of source of infections (e.g. when investigating the source of campylobacteriosis). In contrast, assignment to a single source may be required in other applications such as parentage assignment^77^.

The optimal strategy for selecting loci for humans using either SNPs or microsatellites is S2 (targeting loci with high between-sources allele diversity) while for *Campylobacter* using cgSNPs is S1 (high within-source allele diversity). This difference is due to features within each of the datasets. Based on condition C1 given above that requires high allele diversity between sources, one would naively expect a more accurate attribution when loci with high between-source diversity are targeted (i.e. when using strategy S2). This is indeed the case for source attribution of humans. In contrast, strategy S1 performs marginally better for source attribution of *Campylobacter* isolates. In fact, loci with high within-source diversity in *Campylobacter* genotypes also have high between-source diversity (see Additional file 2: Fig. S6) and are less redundant than those with high between-source diversity (Additional file 2: Fig. S7). For this data set, a high diversity within sources combined with high diversity between sources seems to be a key factor for source attribution. This suggests that a high between-source diversity is necessary in general to distinguish different sources but it is not sufficient to ensure a high-quality source attribution. Based on this and given the formal similarity of the entropy within sources and informativeness^21^ explained in Methods, our results suggest that targeting loci with high informativeness (similar to S2) will not always be optimal compared to S1.

Strategy S3 (reordering loci targeted by strategies S1 and S2) did not bring a significant improvement on S1 or S2 for any of the examples considered here. This suggests that the redundancy of the loci targeted with strategies S1 and S2 does not play an important role in source attribution for these examples. We expect that the relative performance of S3 compared to S1 and S2 will depend on the data set. For instance, S3 could improve on S1 and S2 for data sets with high linkage disequilibrium. For cases in which linkage disequilibrium plays a crucial role, one could devise selection strategies with lower computational complexity than strategy S3. For instance, one could filter out one of the two loci in a pair when such a pair is in high linkage disequilibrium^77^. Strategies focusing on pairs of loci (e.g.^77^) should be computationally faster to apply than S3 but they are expected to be less accurate than S3 in datasets with high linkage disequilibrium.

For *Campylobacter* isolates, we have shown that it is sufficient to use the 10 cgSNPs with the highest within-source entropy to achieve a self-attribution accuracy of ~ 70% that is comparable to that obtained with 25 937 cgSNPs (Fig. 4(a)). In contrast, a much slower increase of the self-attribution accuracy was observed for the human data sets (based on the 659 276 SNPs dataset, one needs more than 1 000 SNPs for the attribution accuracy to saturate). The reason for the slow increase is unclear. It appears there is a lack of loci with high discriminatory power in the human data sets. In fact, loci with high between-source diversity are scarce compared to the *Campylobacter* dataset even in the 659 276 SNPs dataset (compare panels (a) and (b) in Additional file 2: Fig. S6). This difference between human and *Campylobacter* genotypes might be because human SNPs are inherently less diverse than *Campylobacter* SNPs. Another possibility is that 659 276 human SNPs represent a small fraction of the human genome (3.2 GBases) that is perhaps not representative enough in terms of loci diversity (compared to 25 937 cgSNPs which is a larger fraction of the *Campylobacter* genome consisting of 1.8 Mbases). In any case, using 659 276 SNPs is sufficient to achieve highly accurate attribution for humans with the MMD method.

The increase of the self-attribution accuracy with the number of selected loci is also slower for the *P. californicus* and breast cancer examples compared to the *Campylobacter* example (see Additional files 4 and 5). For *P. californicus*, this can be explained by the extremely low between-source diversity of SNPs (Additional file 2: Fig. S6(c)). Due to this, individual SNPs do not efficiently distinguish between the north and south regions in this case even if there is a relatively high within-source diversity (i.e. condition C1 for accurate source attribution is not well satisfied). An accurate distinction between individuals from the north and south regions can only be achieved by combining ~ 100 SNPs; the particular strategy used to select these SNPs does not seem to play a crucial role. Loci diversity is also limited in the breast-cancer proteotypes but the fraction of loci with high between-source diversity is promising. Including more samples in the dataset could potentially enhance the loci diversity in such a way that high attribution accuracy could be achieved by targeting few informative loci. Irrespective of this, it is interesting that even with a limited number of samples, the MMD method already achieves a relatively high self-attribution accuracy using ~ 500 proteomic loci.

The genetic profile of individuals and sources can be represented with a wide range of genetic markers including microsatellites, gene-based markers or SNPs^78^. Data consisting of genotypes which contain large enough sets of highly polymorphic markers will typically offer high discriminatory power. Following this, one can achieve similar attribution accuracies using relatively short multilocus genotypes containing highly polymorphic markers (e.g. microsatellites) or extended genotypes containing less diverse markers (e.g. SNPs). With the current genomic technologies it is becoming increasingly feasible to obtain large sets of SNPs from genomes of many individuals. Combining extended SNP genotypes and fast methods for source attribution such as the MMD provides a significant opportunity for the future of source attribution approaches. Similar arguments apply to other OMIC datasets which are becoming increasingly available, as illustrated in our cancer example.

## Conclusions

The MMD method is very fast, easy to use, suitable for a range of types of loci (e.g. SNP, cgMLST, microsatellite, proteomics loci, etc.) and provides similar assignment accuracies to other methods. The best method for determining the minimum set of loci for optimal attribution varies between datasets. It is therefore prudent to employ a number of methods on each dataset to decide which set of loci are optimal. Some of the locus selection methods can be very computationally intensive (greedy strategies such as S3) and may not be practical to be used in conjunction with current attribution methodologies which are relatively slow. In contrast, the performance of different locus selection strategies can be tested relatively fast with the MMD method. The methods described in this paper are relevant for multiple applications in the life sciences and although they have only been applied to DNA- and proteomics-based methods here, could potentially also be used on other OMIC datasets (e.g. metabolomics) to characterise populations.

## Methods

### *Campylobacter* infectious disease example

Whole genome sequenced *Campylobacter* isolates comprising 500 clinical isolates from human patients and 673 isolates from five food and animal sources were obtained: cattle (150), chicken (150), pig (130), sheep (150) and wild bird (93) (Suppl data file S1). PanSeq^79^ was used to construct a non-redundant pan-genome from all of the 1173 genomes, using a seed genome and identifying regions of ≥ 1000 base pairs (bp) not found in the seed, but present in any other genome at 87% sequence identity cut-off. Loci present in all genomes underwent multiple sequence alignment and were concatenated. This aligned sequence was used to identify SNPs (*n* = 25 937 in the core genome of all isolates (Suppl data file S1)).

### Human evolution example

Assignment of human individuals was illustrated for three data sets with individuals from 7 different geographic regions of the world. The first dataset comprised 5 795 human individuals from 7 different regions (Africa, America, Central/South Asia, East Asia, Europe, Middle East and Oceania). The genotype of each individual was described by 645 microsatellite markers^22^ (Additional file 1: Suppl data file S2). The second dataset comprised 1107 Individuals from the same 7 regions of the microsatellite data set and their genotypes were described by 2 810 SNPs^61^ (Additional file 1: Suppl data file S3). The third dataset comprised 938 humans from the same geographic regions available from the Human Genome Diversity Panel (HGDP). The genotype of each individual was described by 659276 SNPs^60^ (Additional file 1: Suppl data file S4).

### Attribution Methodology

The aim of source attribution is to estimate the probability *p_u,s_* that an individual of unknown origin, *u*, originates from a source *s* from a set 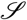 of sources. For haploid genotypes, the unknown individual is characterised by a set of *L* loci, 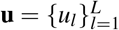. Here, *u_l_* denotes the allele of the individual *u* at locus *l*. The set of possible values taken by the alleles is denoted as 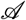. The genetic information of a source *s* is represented by *I_s_* multilocus genotypes; the genotype of an individual *i* in the source *s* is characterised by a set of *L* loci, 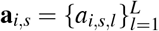. Methods will be described for haploid genotypes but they can be readily extended to diploid genotypes or descriptions of individuals in terms of feature vectors of any kind. In the diploid case, genotypes are characterised by a sequence of *L* loci, each with two alleles: **a**_i,s_ = {(*a*_*i,s,l*_1__). This information can be encoded as a feature vector consisting of 2*L* elements which can be readily used by the MMD method. Alternatively, one can encode the information into a vector of *L* elements by replacing pairs (*a*_*i,s,l*_1__, *a*_*i,s,l*_2__) by a single value, as described in Additional file 1: Suppl file S3 for the 65 533 SNP human genotypes. A method to extract a feature vector from proteomic data is described in Additional file 5.

The source attribution probabilities are summarised by the distribution probability *p_s_* that a randomly chosen individual from a set of *I_u_* isolates of unknown origin (e.g. *I_u_* = 500 *Campylobacter* isolates in Fig. 1) is attributed to the source *s* on average. We assume that *p_s_* has an inherent uncertainty associated with the fact that the set of *I_u_* assigned genotypes is a sample of a larger population of genotypes. In order to estimate the uncertainty of *p_s_*, we estimate its probability distribution by bootstrapping^80,81^ based on the source probabilities 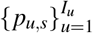 for the *I_u_* assigned genotypes. For a given source, *s*, bootstrapping was implemented as follows: (i) Draw a random sample with replacement of *I_u_* elements from the set 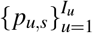. (ii) Calculate the sample mean, 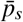, of the selected values of *p_u,s_*. (iii) Repeat steps (i) and (ii) *n_b_* times (*n_b_* = 10^4^ in our calculations). This results in *n_b_* values of 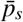 that define our estimate for the distribution of *p_s_*. The error bars in Figs. 1, 2, S1 and S2 correspond to 2.5^th^ and 97.5^th^ percentiles of the *p_s_* distribution.

A Monte-Carlo cross-validation strategy^65,66^was used for self-attribution. More explicitly, *I_u_* individuals were randomly removed from each source population (testing or validation set) and they were attributed to the sources described by the remaining genotypes (learning or training set). The origing of the removed *I_u_* individuals is assumed to be unknown and the probability *p_s_* that any of them is attributed to source *s* (see Figs. 2, S1 and S2) is calculated by bootstrapping, as explained above for source attribution. The self-attribution accuracy is summarised in Figs. 4(a,b) in terms of the total self-attribution probability *p*^sa^ defined as the mean over sources of the probability *p_s_* that individuals from each source are attributed to their source. The confidence interval of *p*^sa^ is estimated by the mean over sources of the 2.5^th^ and 97.5^th^ percentiles of the correct self-attribution probability *p_s_* for each source.

For the *Campylobacter* and humans examples, 50% of the samples were removed from the source to be tested for self-attribution (i.e. *I_u_ = I_s_*/2). Details on the self-attribution analysis for *P. californicus* and breast cancer samples are given in Additional files 4 and 5, respectively.

### The MMD method

The MMD method uses the multilocus genotypes **a**_*u*_ and **a**_*i,s*_ to determine the probability *p_u,s_* as follows:

i. Calculate the Hamming distance^64^, *d*_H_(**u, a**_*i,s*_), between the genotype of unknown origin and genotypes *i* in source *s*.
ii. Obtain a score *σ_u,s_* which quantifies the proximity of *u* to source *s*. The calculation of *σ_u,s_* is based on the cumulative distribution function *F_u,s_*(*λ*) that gives the probability that the Hamming distance between *u* and any genotype of source *s* is smaller than *λ* (see Additional file 2: Fig. S8). The proximity between *u* and each of the sources *s* is measured by the *q*-quantile *λ_u,s_*(*q*) corresponding to the distribution *F_u_*(*λ*). For a given probability *q*, the closest source to *u* is the one with the smallest value of *λ_u,s_*(*q*):

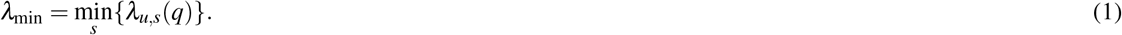 Once *λ*_min_ has been obtained, the score is calculated as *σ_u,s_ = F_u,s_*(*λ*_min_), i.e. it is the probability that the Hamming distance of *u* to any source *s* is *λ*_min_ or smaller. This ensures that sources with high probability to be close to *u* are given a high score (see a graphical representation of the procedure in Additional file 2: Fig. S8).
iii. Estimate the probability that *u* is attributed to source *s* as 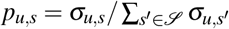. Note that an individual *u* is necessarily attributed to at least one source by the methodology.

The Hamming distance can be calculated in times proportional to the Hamming distance itself^82^. Accordingly, the time complexity for attribution of an individual with MMD is *O*(*d*_max_*I*_Tot_), where *d*_max_ is the maximum Hamming distance between the genotype of unknown origin and the genotypes used to describe sources and 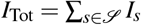 is the total number of genotypes used to describe sources.

The probability *q* is a parameter of the model. In self-attribution tests, the optimal value of this probability was obtained for each source *s* as the value *q*_*_ that maximises the probability *p_s_* that individuals are correctly attributed to their source (in some cases, *q*_*_ can be defined as an interval of *q* where the maximum self-attribution probability is observed). Results of the correct self-attribution probability as a function of *q* are shown in Additional file 2: Figs. S9–S11. The optimal value/interval *q*_*_ depends on the particular set of individuals set as unknown for self-attribution but it is relatively small in all the examples studied here (in most cases, *q*_*_ < 0.1). This makes sense since one would expect that large differences in *σ_u,s_* for different sources would be mainly dictated by few genotypes that are closer to the individual *u* in its source. In particular, setting *q* = 0 defines an extreme version of our algorithm with *λ*_min_ = min_*i,s*_{*d*_H_(**u,a**_*i,s*_)}. In this case, the score *σ_u,s_* is the proportion of genotypes in source *s* that are a distance *d*_min_ from the individual to be assigned, *u*. We checked that self-attribution accuracy is already high when we set *q* = 0 in our examples. In general, however, *σ_u,s_* obeys the extremal value statistics of the Hamming distance for *q* = 0 and might not be reliable enough if the number of genotypes, *I_s_*, used to describe each source, *s*, is not large enough. When *I_s_* is not large enough and extended genotypes are used, individuals of unknown origin tend to be attributed to a single source *s* with probability *p_u,s_* = 1 (i.e. the condition *d*_H_(**u, a**_*i,s*_) = *d*_min_ is only satisfied for one genotype).

For source attribution, *q* cannot be obtained through optimisation since the actual origin of individuals to be attributed is genuinely unknown. In this respect, it can be useful to do self-attribution with genotypes from source populations to estimate a suitable value of *q*. For instance, the source attribution results for human *Campylobacter* isolates shown in Fig. 1(b) correspond to *q* = 0.05 which is the mean of the optimal self-attribution values, *q*_*_, weighted by the number of isolates in each source (see Additional file 2: Fig. S9). In fact, source attribution is not very sensitive to the specific value of *q*, provided it is within the range in which self-attribution probability is high. Compare, for instance, the results for *q* = 0 illustrated in Additional file 2: Fig. S12 with those for *q* = 0.05 in Fig. 1(b).

### The STRUCTURE method

STRUCTURE is a Bayesian clustering model proposed to infer population structure and assign individuals to populations. Following previous works^8,11–13,83^, STRUCTURE was used to estimate *p_u,s_* by setting the number of clusters to be equal to the number of sources (e.g. *K* = 5 for the *Campylobacter* example or *K* = 7 for the humans attribution example). The population structure of the sources was assumed to be know (i.e. we set USEPOPINFO=1 and POPFLAG=1 for the source isolates). In contrast, the population structure of the *I_u_* isolates to be attributed was set as unknown with POPFLAG=0. The results presented are based on runs of 10^4^ MCMC steps following a burn-in period of 10^4^ iterations. The statistics of *p_s_* were obtained from *p_u,s_* as explained above for the MMD method.

### The ADMIXTURE method

ADMIXTURE uses multilocus genotype data for efficient estimation of ancestry of unrelated individuals^40^. ADMIXTURE infers the ancestry of individuals in terms of the admixture proportion *h_u,s_* of the genome of individual *u* that originated from population *s*. In the supervised version of ADMIXTURE^48^, the ancestry of reference populations is determined by the genotypes of the individuals in the training set (i.e. all individuals except the *I_u_* individuals selected for the validation set). The admixture proportion *h_u,s_* for individuals in the validation set quantifies the proportion of their genotype originating from the reference population *s*. This can be regarded as a measure of genetic proximity between the individual *u* and population *s*, formally similar to the attribution probability *p_u,s_* estimated by the MMD method. Following this, our application of ADMIXTURE for source attribution uses a supervised analysis and assumes that the attribution probability is *p_u,s_* = *h_u,s_*.

### Information theory: Loci diversity and redundancy

We quantify the allele diversity in terms of the Shannon entropy, a measure of the information (in bits) necessary to describe the uncertainty of random variables^57^. The Shannon entropy is increasingly used as a diversity index in ecology^84,85^ and population genetics^86–88^. In our application, the random variables are the alleles found in the genotypes of sources at a locus *l*. More explicitly, we consider the probability *π_a,l,s_* that an allele takes the value *a* at the locus *l* and 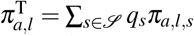 which gives the allele probability pooled over sources. Here, 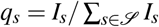 is the proportional weight of each source. The total allele diversity in a locus *l* is quantified by the Shannon entropy of the distribution 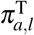,

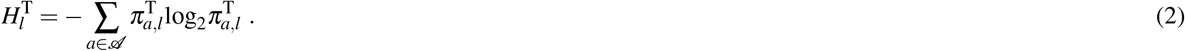

For example, the fact that the maximum number of alleles in a SNP is 4 (A, T, C and G) implies that the entropy 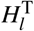 could take any value between 0 (the same allele in all genotypes) and 2 bits (maximal diversity when each allele appears in 1/4 of the genotypes). As expected for any measure of allele diversity, the larger the number of alleles in a locus, the larger the Shannon entropy. Microsatellites^22^ or gene-based markers^24^ are characterised by larger sets 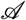 of possible alleles and can be more diverse than SNPs, i.e. they have larger values of 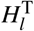 which are mostly associated with a larger contribution of the diversity within-sources, 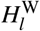 (see Additional file 2: Fig. S6).

The entropy 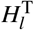 gives the allele diversity for subtypes in *all* the sources. The condition C1 given above for selection of informative markers, however, suggests that it is the allele diversity between sources the one that could play a major role on the assignment power of loci. Accordingly, we split the total entropy 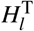 in two contributions^84,85^: One accounting for the diversity *within* sources,

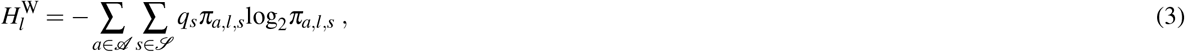

and another measuring the diversity *between* sources,

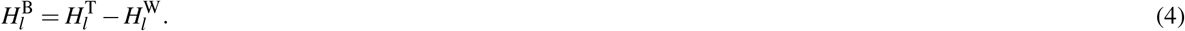

Basic algebraic manipulations show that 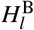 is formally similar to the informativeness introduced in^21^. Our interpretation of 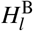 is, however, slightly different to that proposed in^21^ since we derived it as an index to distinguish sources rather than as a measure of the information gained when adding new loci to the selection used for attribution.

### Mutual information and redundancy of loci

The source attribution discriminatory power of a set of *n* loci is typically not *n* times larger than the discriminatory power of each isolated locus. This is due to the fact that loci are not statistically independent, i.e. there is some redundant information when considering several loci. The concept of loci redundancy was used in this work with two aims: To select pairs of loci with low redundancy in strategy S3 and to assess the extent to which strategies S1 and S2 satisfy the condition C2 of low redundancy.

The elementary quantity in our estimates of loci redundancy is the mutual information between pairs of loci. Given a pair of loci, (*l, l*′), it is defined as^57,87,88^:

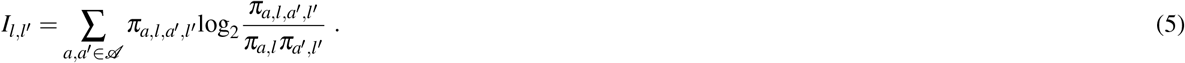

Here, *π_a,l,a′,l′_* is the joint probability distribution for alleles in locus *l* and *l*′. Within the context of population genetics, *I_l,l′_* has been used to quantify the linkage disequilibrium between loci *l* and *l*′^87^. The mutual information takes values 0 ≤ *I_l,l′_* ≤ min{*H_l_, H_l_*′}. In particular, it is null, i.e. *I_l,l′_* 0, when the allele distributions of the two loci are independent. In general, *I_l,l′_ ≤ I_l,l_* meaning that a locus contains as much information about itself as any other locus can provide. In other words, a locus *l* is maximally redundant with respect to itself. In the case *l = l*′, the mutual information coincides with the Shannon entropy, *H_l_ = I_l,l_*. Within the context of this work, mutual information is used as a measure of the linkage disequilibrium between the pairs of loci *l* and *l*′. In fact, *I_l,l′_* is proportional to the widely-used^56^ measure for linkage disequilibrium, *r*^2^, in the limit of *π_a,l,a′,l′_* ≤ 2*π_a,l_π_a′l′_*^87,89^. Despite the similarity with classical measures such as *r*^2^, the mutual information gives an intuitive interpretation of linkage disequilibrium and, as discussed in^87,89,90^, has some other advantages over classical measures.

For our particular application, *I_l,l′_* allows us to naturally define a measure of loci redundancy relevant to strategy S3. The redundancy *R_n_* of the *n*-th locus added to a list of *n* − 1 previously selected loci is given by the following formula:

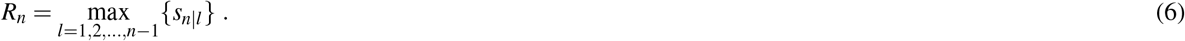

Here, 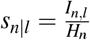 is the reduction in uncertainty of a locus *l* = 1,2,…,*n* − 1 when the *n*–th locus is added to the set used for source attribution. The definition of *s_n|l_* and the whole redundancy analysis is restricted to loci with *H_n_* > 0; loci with *H_n_* = 0 consist of a single allele and are excluded from the analysis since they have a null discriminatory power. From Eq. (6), *R_n_* can be interpreted as the maximal reduction in uncertainty achieved when adding the n-th locus to the list of selected loci. By definition, 0 ≤ *R_n_* ≤ 1. The case *R_n_* = 0 corresponds to the smallest possible redundancy of locus *n* and is observed when the allele distribution at such locus is statistically independent of the allele distribution at any of the previously selected loci *l*. The case *R_n_* = 1 indicates that *s_n|l_* = 1 for at least one of the previously selected loci, thus indicating that the allele distribution of the newly added locus, *n*, is identical to the allele distribution of at least one of those that were previously selected. In this case, the locus *n* would not contribute to enhance the discriminatory power of the set of selected loci.

## Acknowledgements (not compulsory)

The *Campylobacter* work in this project was supported by Food Standards Scotland project FSS00017 and the Scottish Government (Rural and Environment Science and Analytical Services Division) project A13559368.

## Author contributions statement

FPR, KF and NS planned the project and designed the research. FPR with OR and BSL wrote the software, retrieved the data and performed the experiments. FPR wrote the manuscript with NS and KF. All authors reviewed the manuscript and approved the final version of it.

## Additional information

### Availability of data and materials

Data used in this work are available from https://figshare.com/s/726d493387b501c4b70a. An executable version of the MMD program can be downloaded from https://figshare.com/s/125861c0a0499ff3101b. The software is based on the MMD R package developed in this project.

## Additional file 1: Supplementary data files

The data sets for the six examples studied in this work are available as compressed ZIP files from https://figshare.com/s/726d493387b501c4b70a

For each example, the corresponding ZIP file contains two text files: A file with extension *.pop which lists the population corresponding to each genotype and a file with the same name but in comma-separated values (csv) format which contains the genotypes (or proteotype for the breast cancer example). *Loci must be in integer format* for our implementation of the MMD method in R.

### Supplementary data file S1, Campylobacter_25 937SNP.zip - *Campylobacter* data

This ZIP file contains the data files Campylobacter_25937SNP.csv and Campylobacter_25937SNP.pop which give information on 1173 *Campylobacter* isolates. The Campylobacter_25937SNP.csv file contains genotypes consisting of 25 937 cgSNPs for the 1173 *Campylobacter* isolates. Each row in the file gives the genotype of one isolate with the following format:

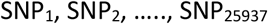

SNPs can take values 1, 2, 3, 4 which correspond to the nucleotides A, T, G and C. Missing loci in the list of SNPs are coded by a negative number that is different for each row (i.e. different for each genotype). In this way, missing loci contribute to an increase of the Hamming distance between pairs of genotypes even if they are missing from both genotypes.

The file Campylobacter_25 937SNP.pop consists of 1173 lines specifying the host name for each *Campylobacter* isolate. The host names are Human, Cattle, Chicken, Pig, Sheep and WB.

### Supplementary data file S2, Human_645microsatellite.zip - Human microsatellite data

This dataset is an adaptation of the data used in Ref. [1] to the format needed for the MMD software. The ZIP file contains the files Human_645microsatellite.csv and Human_645microsatellite.pop. The file Human_645microsatellite.csv contains genotypes with 645 microsatellites for 5795 human individuals. Each row gives the genotype of one individual with the following format:

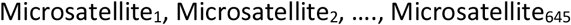

Different integer numbers codes different microsatellites. As in Suppl. data file S1, missing alleles are indicated by a different negative number for each row.

The file Human_645microsatellite.pop lists the region of each individual. The names of the regions are AFRICA, AMERICA, CENTRAL_SOUTH_ASIA, EAST_ASIA, EUROPE, MIDDLE_EAST and OCEANIA.

### Supplementary data file S3, Human_2810SNP.zip - Human 2810 SNP data

This dataset is an adaptation of the data used in Ref. [2] to the format needed for the MMD software. The ZIP file contains two files: Human_2810SNP.csv and Human_2810SNP.pop. The Human_2810SNP.csv file contains diploid genotypes with 5620 loci for 1107 human individuals. Each row gives the genotype of one individual with the following format:

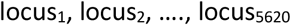

Loci can take values 1, 2, 3, 4 which correspond to the nucleotides A, T, G and C. Each locus gives the allele of one of the copies in a pair of alleles (therefore, there are twice as many loci as SNPs). Missing loci are coded by a negative number that is different for each row (i.e. different for each genotype).

The file Human_2810SNP.pop lists the region of each individual. The names of the regions are AFRICA, AMERICA, CENTRAL_SOUTH_ASIA, EAST_ASIA, EUROPE, MIDDLE_EAST and OCEANIA.

### Supplementary data file S4, Human_659276SNP.zip - Human 659 276 SNP data

This dataset is an adaptation of the data used in Ref. [3] to the format needed for the MMD software. The ZIP file contains three files: Human_659276SNP.csv and Human_659276SNP.pop and Population_names_938.pop. The Human_659276SNP.csv file contains genotypes with 659276 diploid SNPs for 938 human individuals. Each row gives the genotype of one individual with the following format:

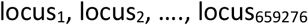

Loci take values {1,2,…,10} which represent pairs of alleles. The conversion is given by the following table:

| AA | AT, TA | AG, GA | AC, CA | TT | TG, GT | TC, CT | GG | GC, CG | CC |
| --- | --- | --- | --- | --- | --- | --- | --- | --- | --- |
| 1 | 8 | 5 | 9 | 3 | 7 | 6 | 2 | 10 | 4 |

Missing loci are coded by a negative integer that is different for each individual. Replacing pairs of alleles by a single number as indicated in the table leads to genotypes of length equal to the number of SNPs. We checked that source attribution based on the coding of pairs of alleles is as accurate as that achieved by coding each individual copy A, T, G, C by 1, 2, 3, 4 (the later coding leads to genotypes of 1 318 552 loci which can also be handled by the MMD program).

The file Human_659276SNP.pop lists the region of each individual. The names of the regions are AFRICA, AMERICA, CENTRAL_SOUTH_ASIA, EAST_ASIA, EUROPE, MIDDLE_EAST and OCEANIA. The file Population_names_938.pop lists the population of origin of each individual. There are 53 populations in total.

### Supplementary data file S5, Pcalifornicus_3699SNP.zip - Giant Californian sea cucumber *Parastichopus californicus* data

This dataset is an adaptation of the data used in Ref. [4] to the format needed for the MMD software. The ZIP file contains three files: Pcalifornicus_3699SNP.csv and Pcalifornicus_3699SNP.pop. The Pcalifornicus_3699SNP.csv file contains genotypes with 7398 loci for 717 human individuals. Each row gives the genotype of one individual with the following format:

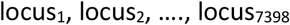

Loci can take values 1, 2, 3, 4 which correspond to the nucleotides A, T, G and C. Each locus gives the allele of one of the copies in a pair of alleles (therefore, there are twice as many loci as SNPs). Missing loci are coded by a negative number that is different for each row (i.e. different for each genotype). The file Pcalifornicus_3699SNP.pop lists the region (North or South) of each individual.

### Supplementary data file S6, Breast_Cancer_proteome.zip – Breast cancer proteomic data

This dataset is an adaptation of the proteomic data used in Ref. [5] to a format suitable for the MMD software. The ZIP file contains three files: Breast_Cancer_proteome.csv and Breast_Cancer_proteome.pop. The Breast_Cancer_proteome.csv file contains genotypes with 65533 loci for 40 human individuals. Each row gives the genotype of each of the samples with the following format:

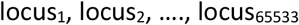

Loci can take values 0 or 1, corresponding to zero or positive values for the mass spectrum intensity, respectively. The Breast_Cancer_proteome.pop file list the cancer subtype of each sample. Subtypes are ERPR, Her2 and TN.

## Additional file 2: Supplementary figures

**Fig S1.**
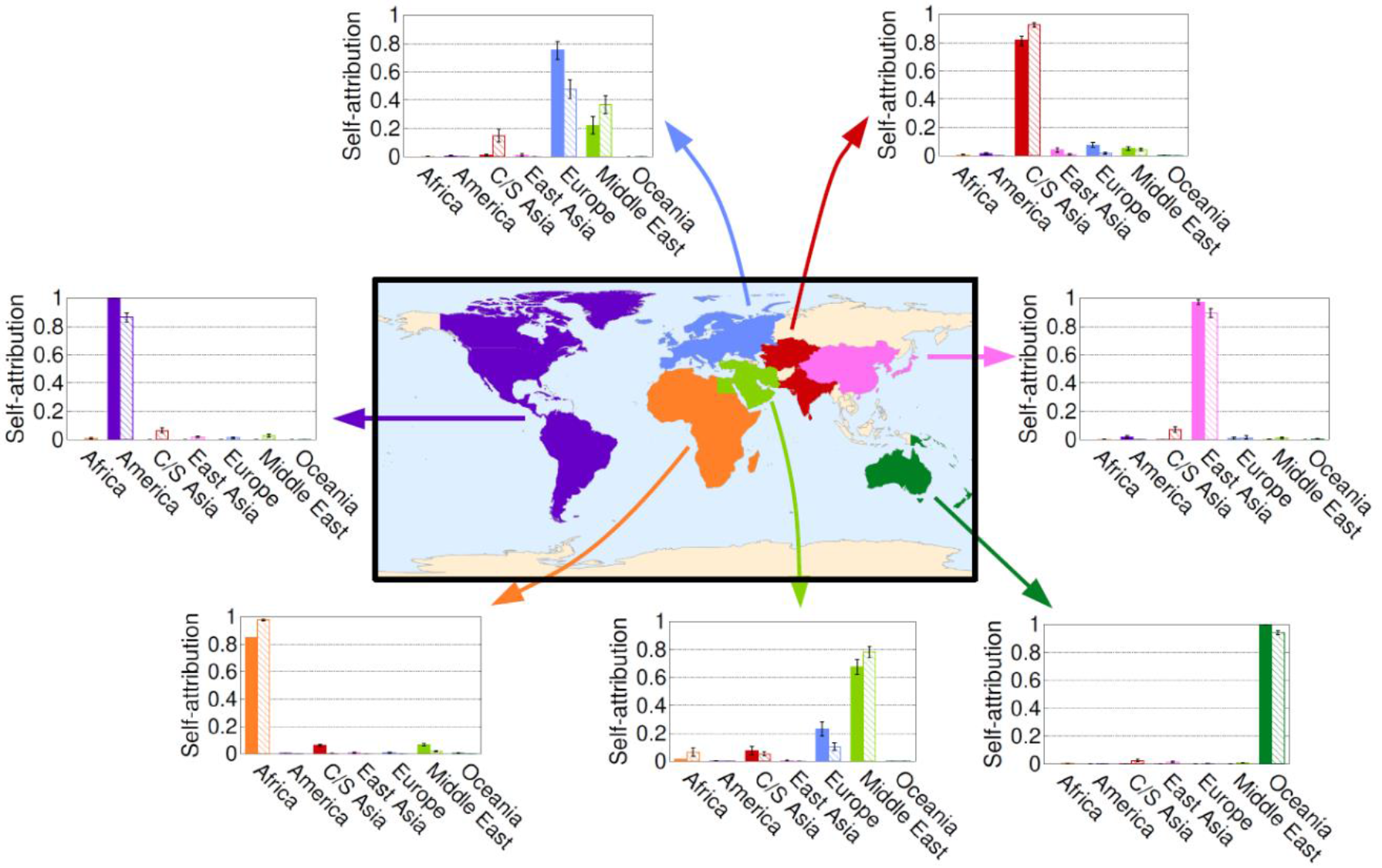
Self-attribution of humans characterised by 645 microsatellite genotypes. Bar charts show the probability distribution *p_s_*. For a given source (region) *s, p_s_* gives the probability that any individual from the region indicated in the map is attributed to *s*. Solid and hatched bars show the results obtained with the MMD method and STRUCTURE, respectively.

**Fig S2.**
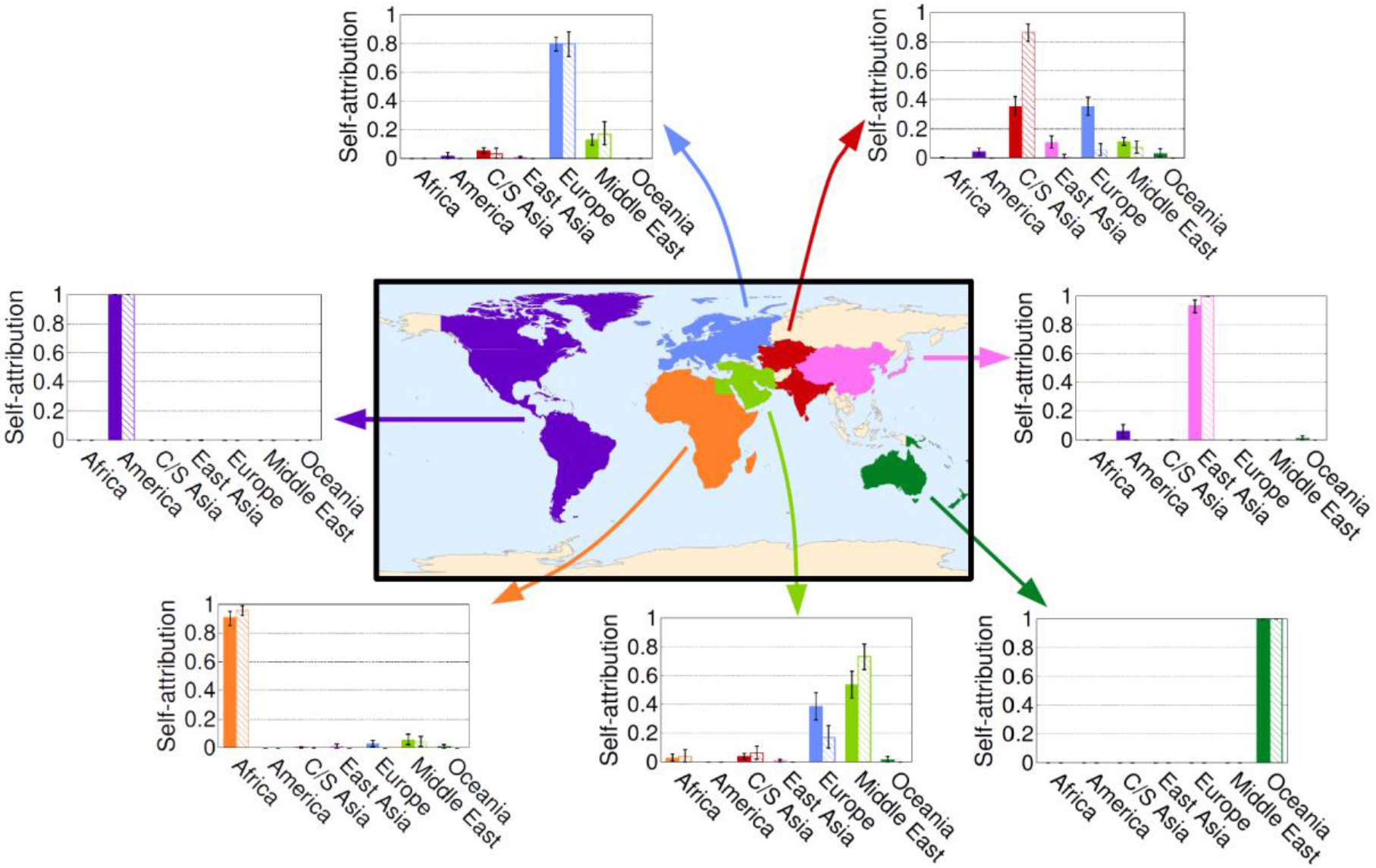
Self-attribution of humans characterised by 2 810 SNP genotypes. Bar charts show the probability distribution *p_s_*. For a given source (region) *s, p_s_* gives the probability that any individual from the region indicated in the map is attributed to *s*. Solid and hatched bars show the results obtained with the MMD method and STRUCTURE, respectively.

**Fig S3.**
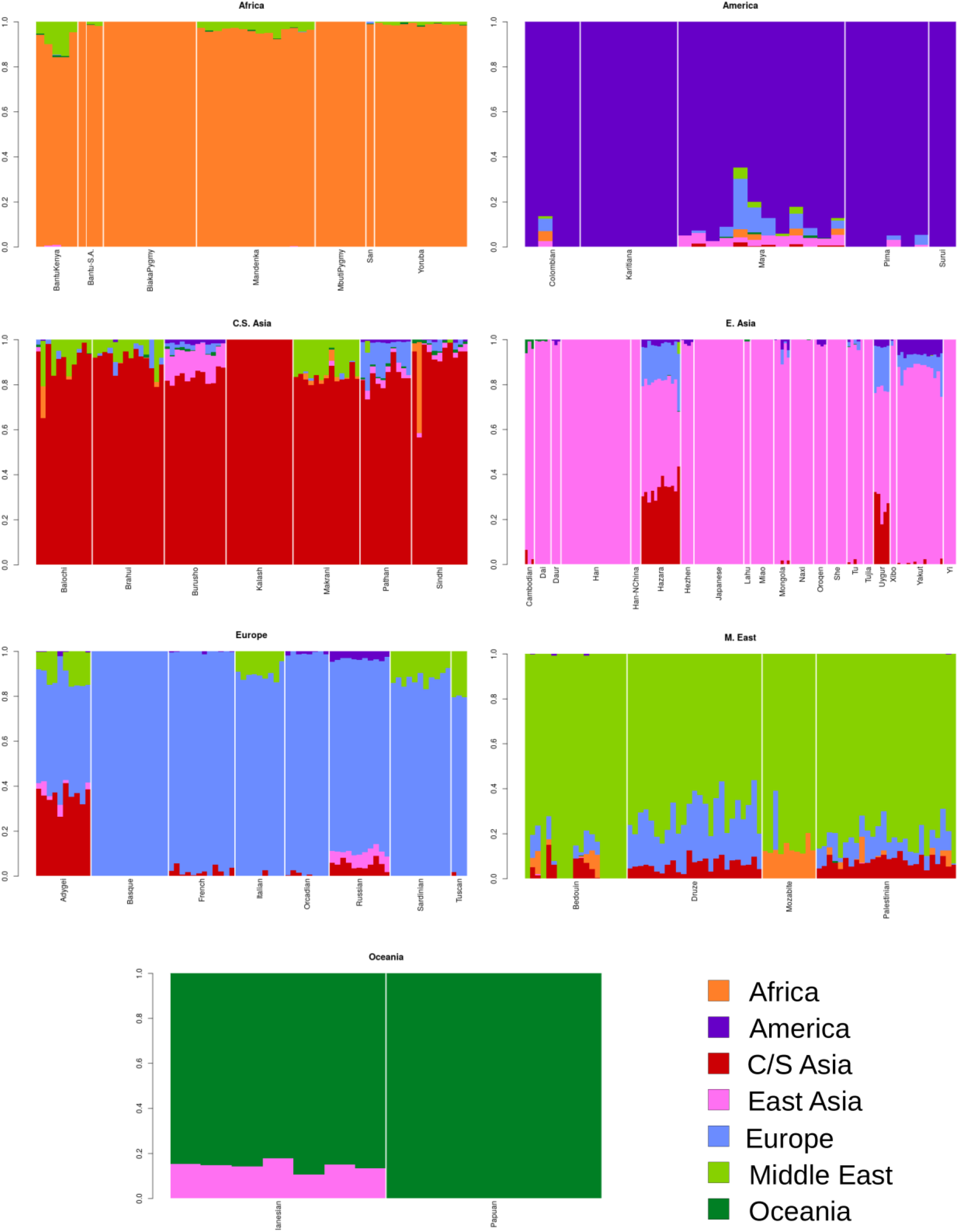
Supervised ADMIXTURE analysis of ancestry based on 659 271 SNP genotypes. Each panel shows the inferred admixture of 50% of individuals *u* selected from the geographical region indicated in the title of the panel. Each individual *u* is indicated by a vertical line, which is partitioned in segments of different colours that represent the admixture proportion *h_u,s_* of the individual from region *s*. The correspondence between regions and colours is given by the legend. Vertical white lines separate individuals from different populations.

**Fig S4.**
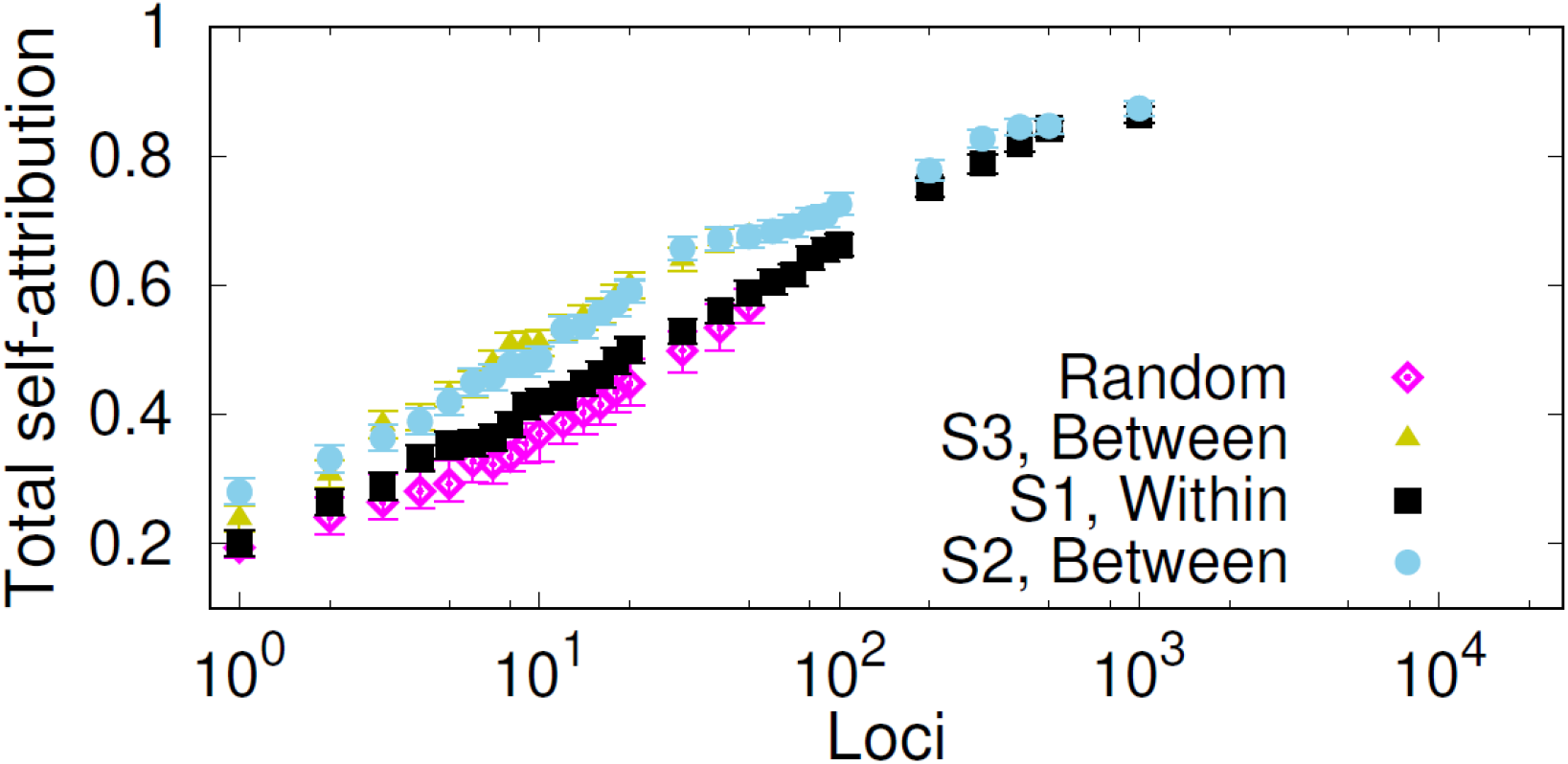
Selection of markers for self-attribution of humans based on 645 microsatellite genotypes. Symbols show the self-attribution probability *p*^sa^ that individuals from any of the 7 regions in the data set are correctly attributed to their region. The probability is plotted as a function of the number of SNPs selected at random and with strategies S1 (loci ranked in decreasing within-source diversity), S2 (loci ranked in decreasing between-source diversity) and S3 (reordering the loci ranking of S2 to reduce loci redundancy).

**Fig S5.**
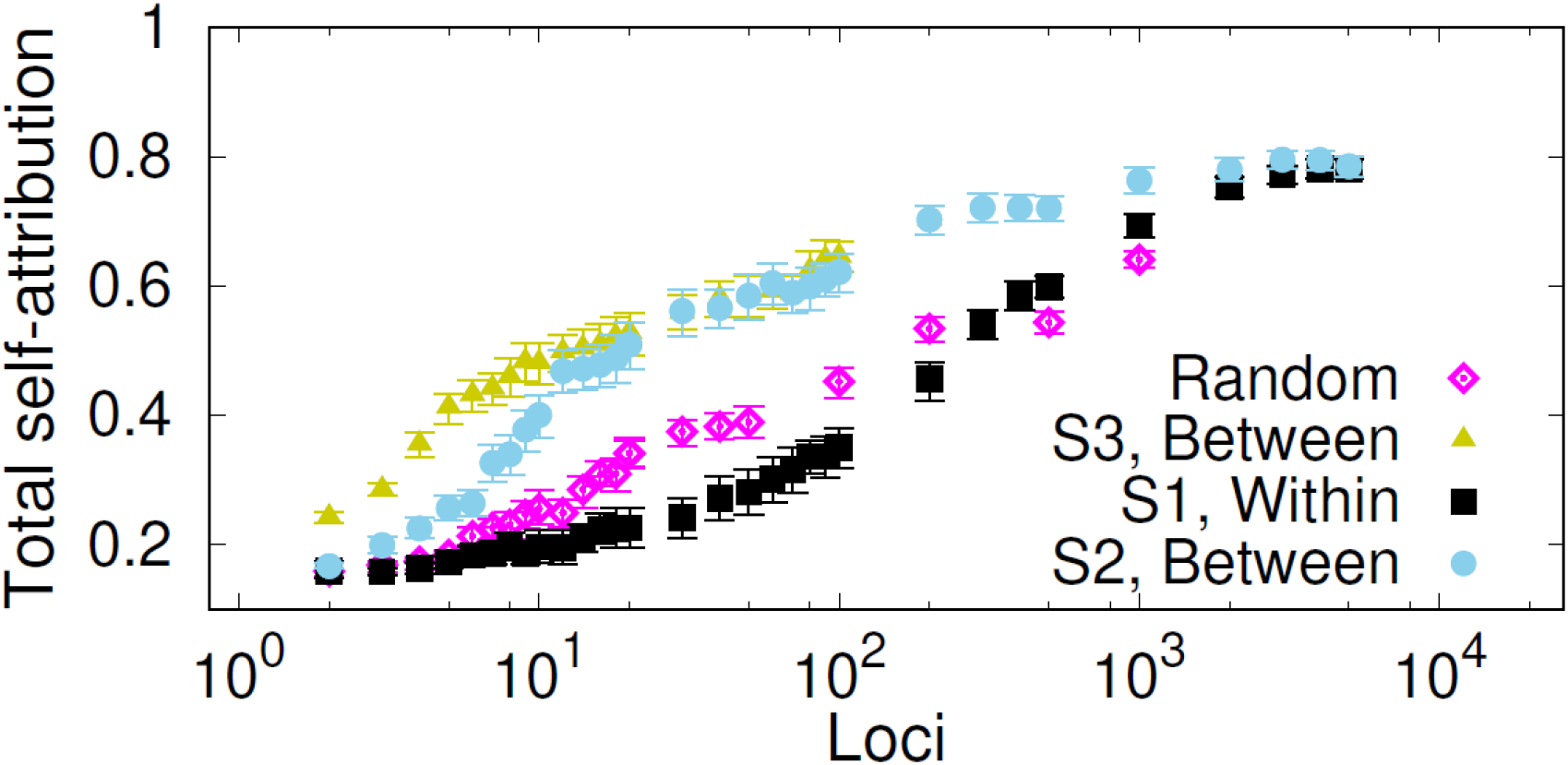
Selection of markers for self-attribution of humans based on 2 810 SNP genotypes. Symbols show the self-attribution probability *p*^sa^ that individuals from any of the 7 regions in the data set are correctly attributed to their region. The probability is plotted as a function of the number of SNPs selected at random and with strategies S1 (loci ranked in decreasing within-source diversity), S2 (loci ranked in decreasing between-source diversity) and S3 (reordering the loci ranking of S2 to reduce loci redundancy).

**Fig S6.**
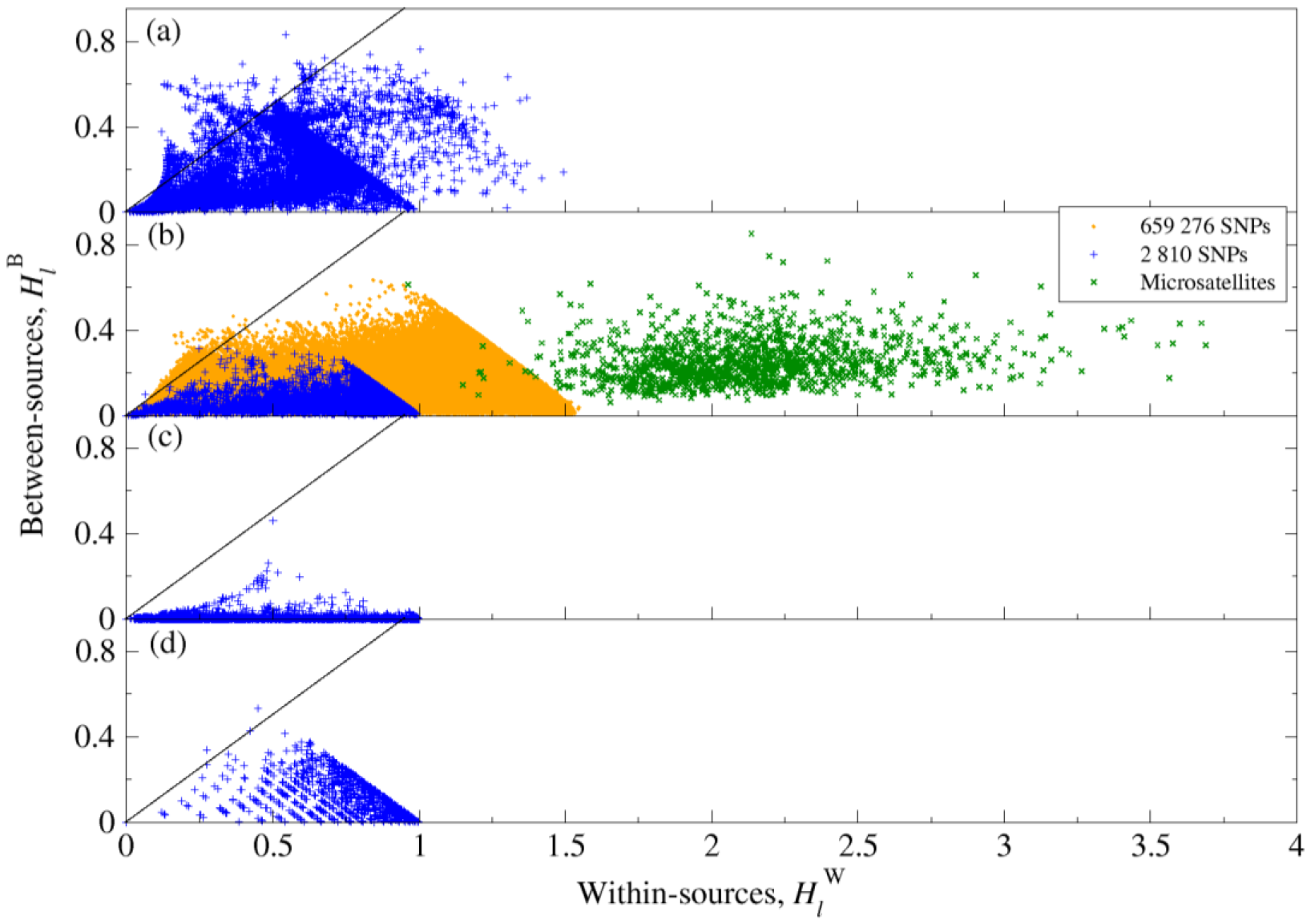
Within- and between-sources allele diversity quantified by entropies. Symbols show the entropy between sources, 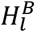, vs. the entropy within sources, 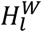, for (a) *Campylobacter* cgSNPs, (b) Human SNPs and microsatellites, (c) *P. californicus* SNPs and (d) breast cancer proteotypes. Most of the data lay on the right of the line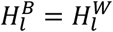, i.e. the diversity within sources is larger than the entropy between sources for most loci in all the data sets.

**Fig S7.**
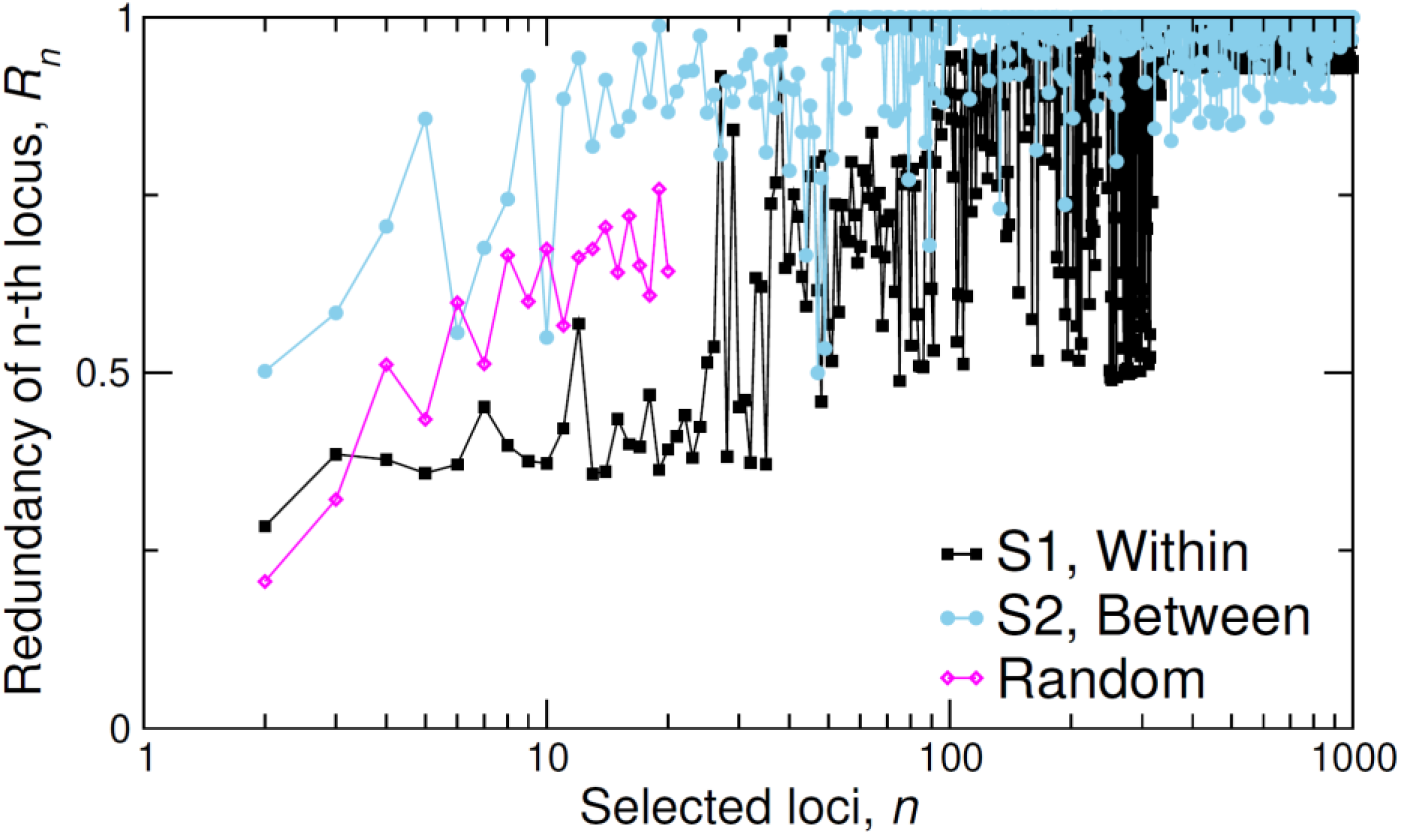
Redundancy *R_n_* of the *n*-th selected locus from 25938 cgSNP *Campylobacter* genotypes. The redundancy *R_n_* is given by Eq. (6) in the main text. The dependence of *R_n_* on *n* is shown for randomly selected loci (pink diamonds) and loci ranked with strategies S1 (black squares) and S2 (blue circles).

**Fig S8.**
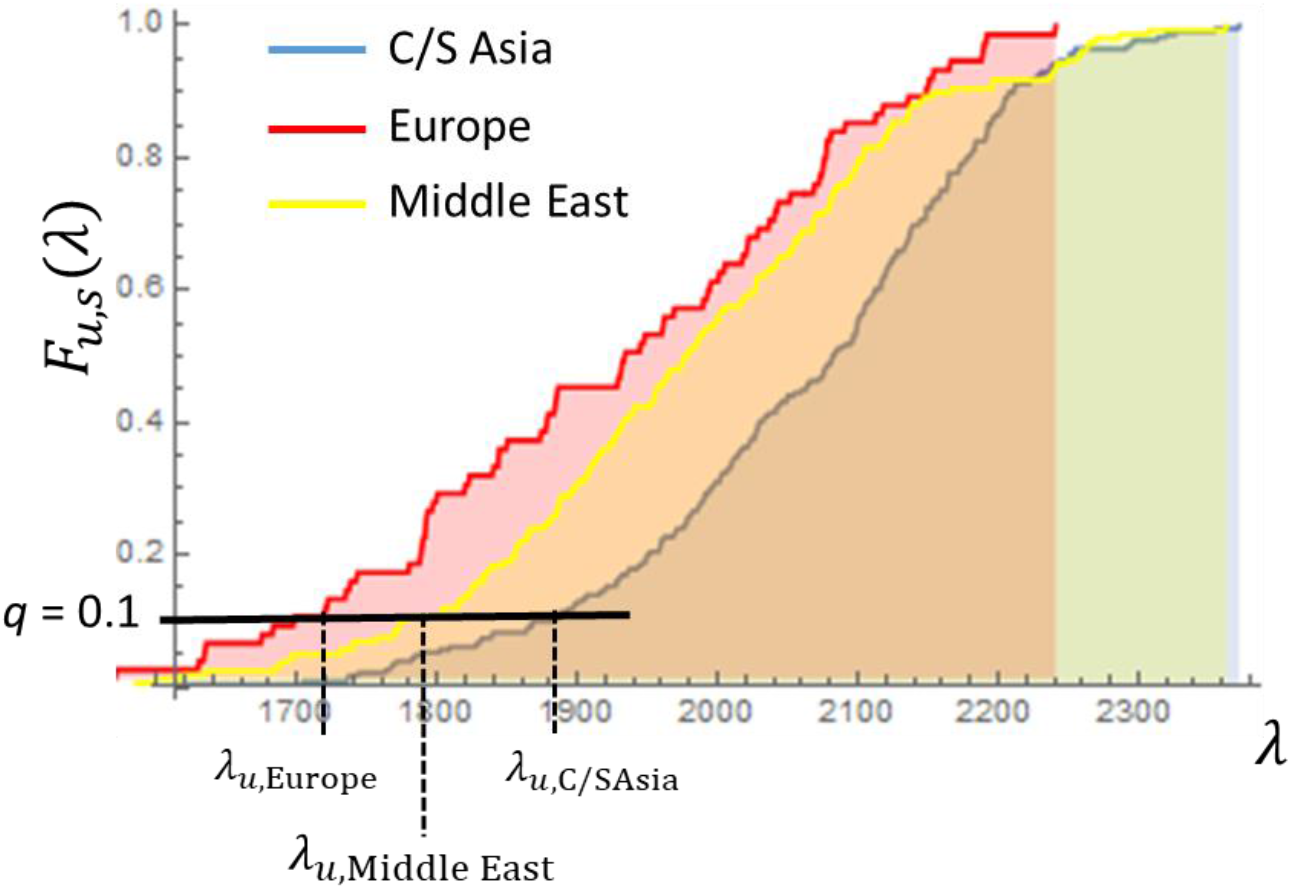
Example of the determination of the minimum *q*-quantile, *λ*_min_(*q*), in the MMD method. The curves show the cumulative distribution function *F_u,s_*(*λ*) which gives the probability that the Hamming distance between an individual of unknown origin, *u*, and any genotype from source *s* is smaller than *λ*. This example corresponds to 2810 SNP genotypes of humans from three regions: C/S Asia, Europe and Middle East. For a probability *q* = 0.1, one obtains the minimum *q*-quantile*λ_min_*(q) = *λ_u,Europe_*. The genotype of *u* is closest to Europe, followed by Middle East and C/S Asia.

**Fig S9.**
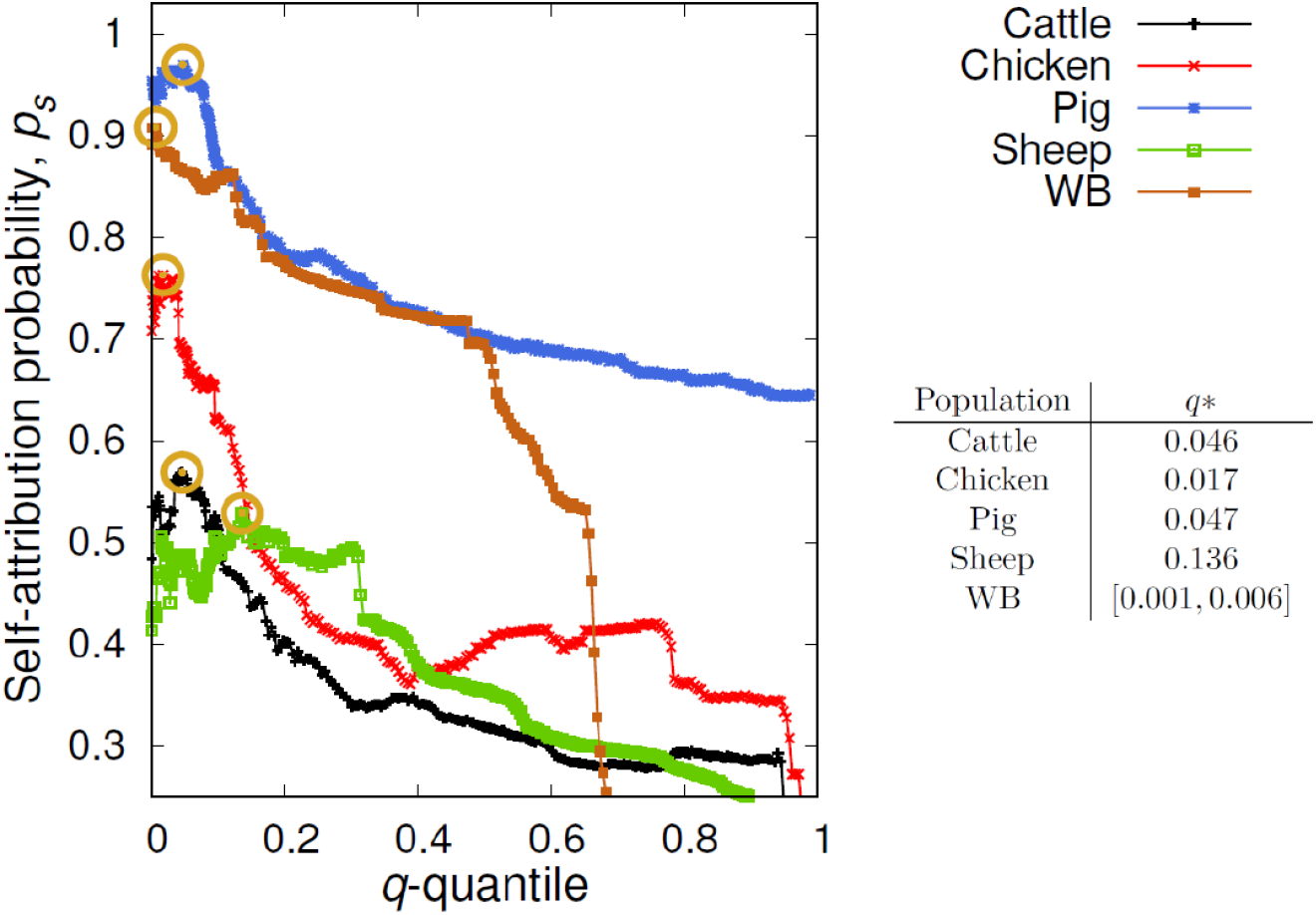
Self-attribution probability *p_s_* based on 25938 cgSNP *Campylobacter* genotypes. Different curves show the probability *p_s_* that a removed individual from a food reservoir *s* (see legend) is correctly attributed to *s*. The circles indicate the point with maximum self-attribution probabilitiy for each source population. The values *q*_*_ (or intervals) of the q-quantile giving the maximum probability are given in the table.

**Fig S10.**
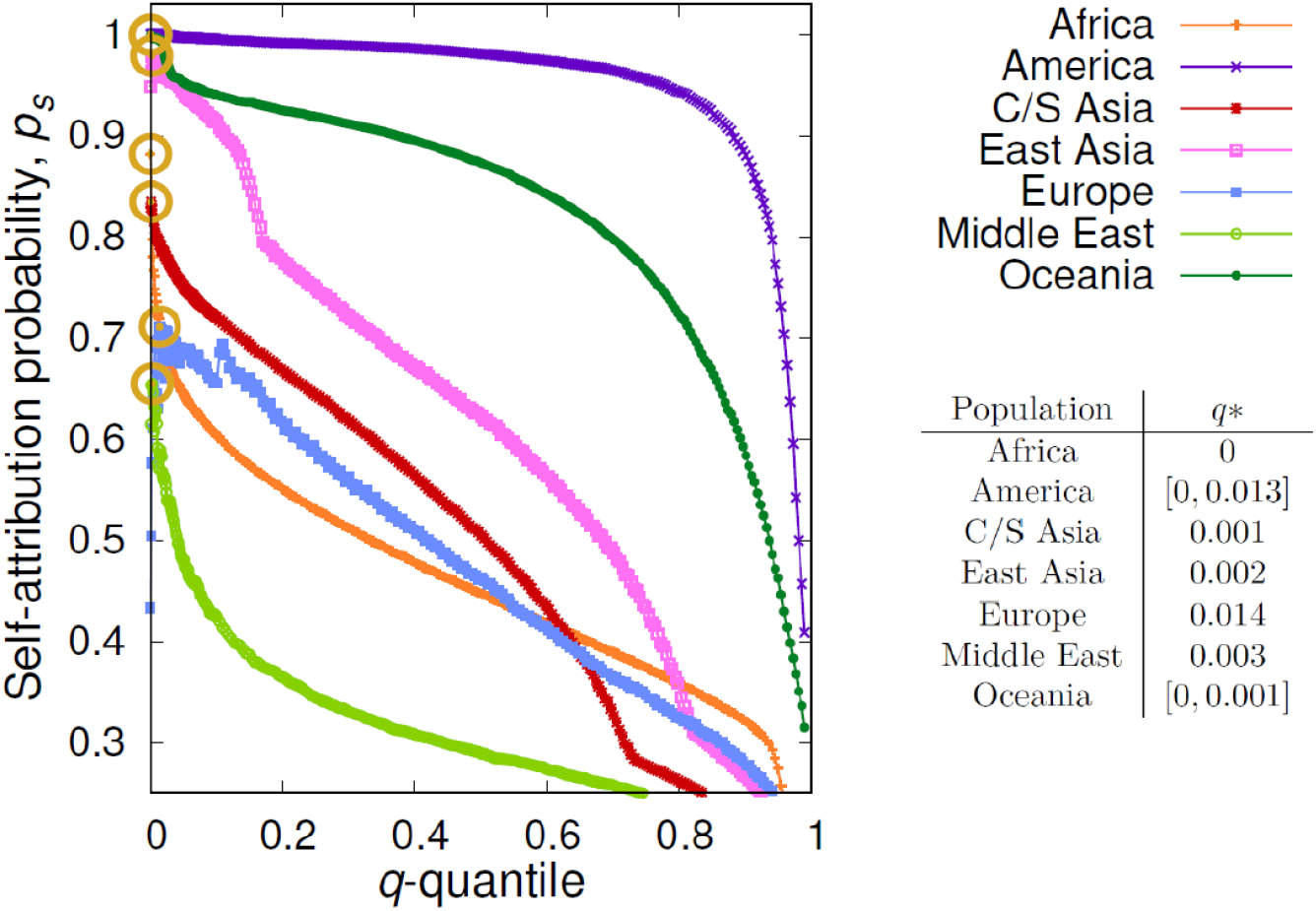
Self-attribution probability *p_s_* based on 645 microsatellite human genotypes. Different curves show the probability *p_s_* that a removed individual from region *s* (see legend) is correctly attributed to *s*. The circles indicate the point with maximum self-attribution probabilitiy for each source population. The values *q*_*_ (or intervals) of the q-quantile giving the maximum probability are given in the table.

**Fig S11.**
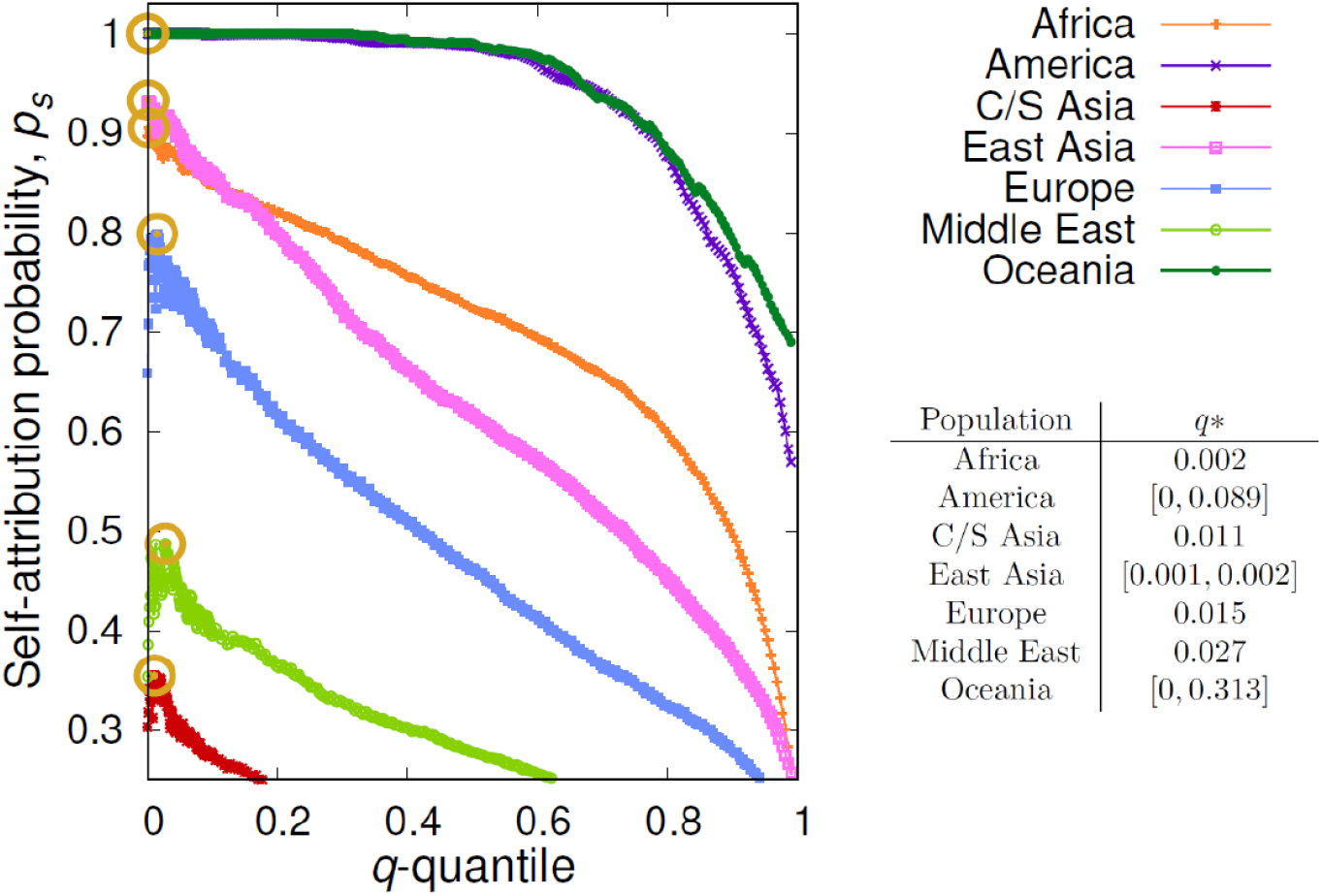
Self-attribution probability *p_s_* based on 2 810 SNP human genotypes. Different curves show the probability *p_s_* that a randomly individual from region *s* (see legend) is correctly attributed to *s*. The circles indicate the point with maximum self-attribution probabilitiy for each source population. The values *q*_*_ (or intervals) of the q-quantile giving the maximum probability are given in the table.

**Fig S12.**
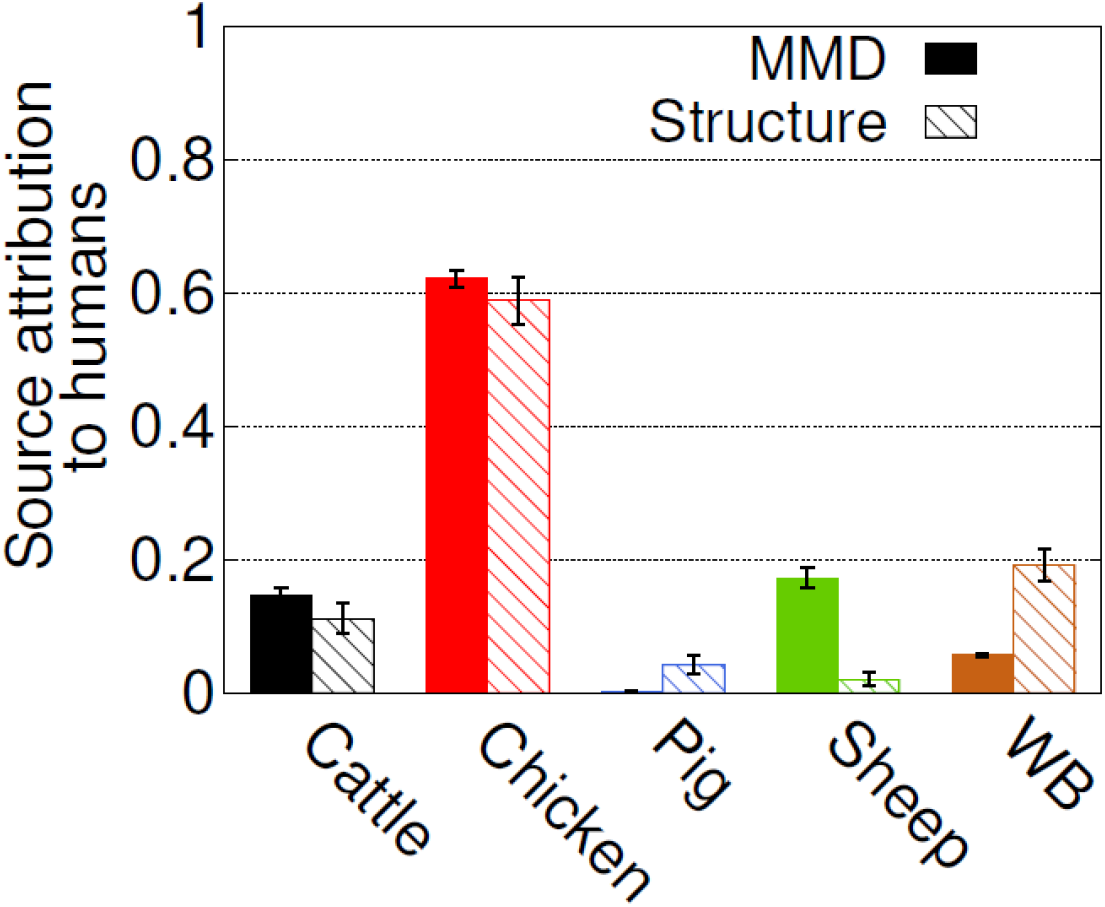
Source attribution of human 500 human Campylobacter isolates. The bar chart shows the source attribution probability distribution ps obtained with MMD (quantile q=0, solid bars) and STRUCTURE (hatched bars) methods. Results based on 25938 cgSNP genotypes.

**Fig S13.**
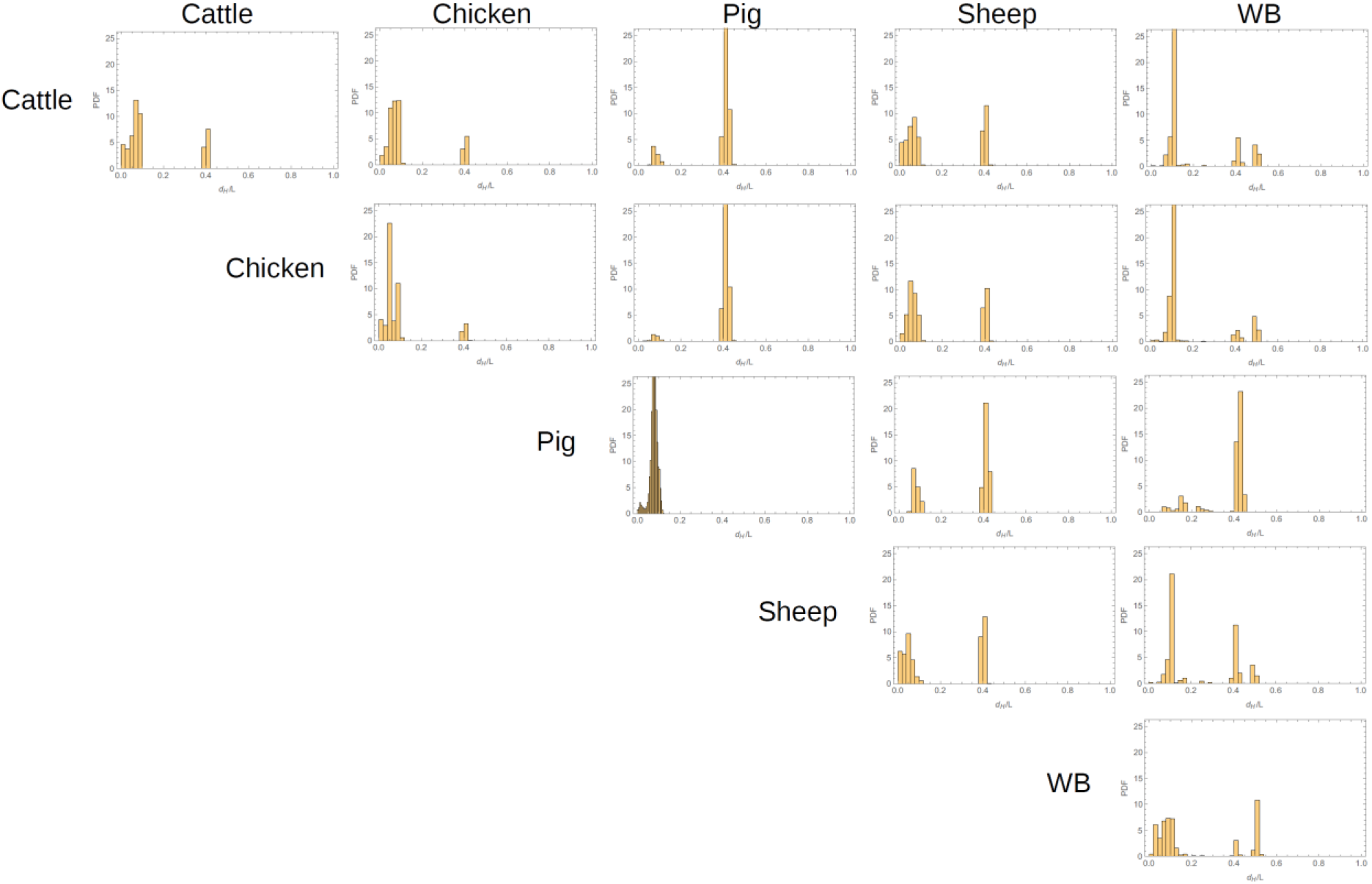
Hamming distance between and within sources for *Campylobacter* isolates based on 25 938 cgSNP genotypes. Each panel shows the density histogram for the Hamming distance, *d_H_*, between pairs of genotypes from the sources indicated by the row and column labels. The horizontal axis of each plot ranges between 0 and 1 and shows the Hamming distance normalised to the total number of loci, *L* = 25938, in each genotype.

## Additional file 3: Supplementary tables

**Table S1.** Allele-frequency divergence for *Campylobacter* sources based on 25938 cgSNPs. Cells indicate the value of the allele-frequency divergence and blue bars give a visual representation of the value in the cell.

|  | Cattle | Chicken | Pig | Sheep |
| --- | --- | --- | --- | --- |
| WB | 0.0645 | 0.0959 | 0.3015 | 0.0828 |
| Sheep | 0.0236 | 0.0583 | 0.222 |  |
| Pig | 0.3279 | 0.4092 |  |  |
| Chicken | 0.0292 |  |  |  |

**Table S2.** Allele-frequency divergence for Humans based on 645 microsatellite genotypes. Cells indicate the value of the allele-frequency divergence and blue bars give a visual representation of the value in the cell.

|  | Africa | America | C/S Asia | East-Asia | Europe | Middle-East |
| --- | --- | --- | --- | --- | --- | --- |
| Oceania | 0.0412 | 0.0506 | 0.0264 | 0.0248 | 0.0349 | 0.033 |
| Middle-East | 0.0185 | 0.0423 | 0.0079 | 0.024 | 0.0044 |  |
| Europe | 0.0261 | 0.0412 | 0.0081 | 0.0243 |  |  |
| East-Asia | 0.0357 | 0.0316 | 0.017 |  |  |  |
| C/S Asia | 0.0244 | 0.035 |  |  |  |  |
| America | 0.0548 |  |  |  |  |  |

**Table S3.** Allele-frequency divergence for Humans based on 2 810 SNP genotypes. Cells indicate the value of the allele-frequency divergence and blue bars give a visual representation of the value in the cell.

|  | Africa | America | C/S Asia | East-Asia | Europe | Middle-East |
| --- | --- | --- | --- | --- | --- | --- |
| Oceania | 0.0548 | 0.0656 | 0.0404 | 0.0398 | 0.0515 | 0.044 |
| Middle-East | 0.0306 | 0.0438 | 0.0054 | 0.0306 | 0.005 |  |
| Europe | 0.047 | 0.0488 | 0.0073 | 0.0374 |  |  |
| East-Asia | 0.0482 | 0.0344 | 0.0239 |  |  |  |
| C/S Asia | 0.0347 | 0.0377 |  |  |  |  |
| America | 0.0663 |  |  |  |  |  |

**Table S4.** Allele-frequency divergence for Humans based on 659 276 SNP genotypes. Cells indicate the value of the allele-frequency divergence and blue bars give a visual representation of the value in the cell.

|  | Africa | America | C/S Asia | East-Asia | Europe | Middle-East |
| --- | --- | --- | --- | --- | --- | --- |
| Oceania | 0.064121636 | 0.00901646 | 0.009537095 | 0.00397227 | 0.027877831 | 0.019187964 |
| Middle-East | 0.067584773 | 0.018132307 | 0.002310077 | 0.012970799 | 0.002605231 |  |
| Europe | 0.088160012 | 0.025433483 | 0.00625519 | 0.020048184 |  |  |
| East-Asia | 0.064896711 | 0.00263804 | 0.005942497 |  |  |  |
| C/S Asia | 0.064709747 | 0.010672819 |  |  |  |  |
| America | 0.068563086 |  |  |  |  |  |

**Table S5.**
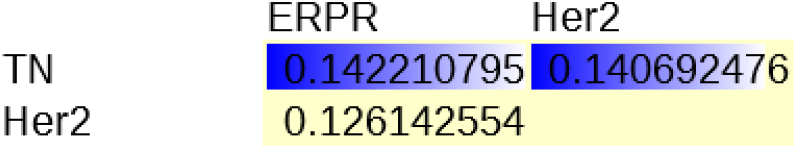
Allele-frequency divergence for breast tumour proteotypes. Cells indicate the value of the allele-frequency divergence and blue bars give a visual representation of the value in the cell.

|  | ERPR | Her2 |
| --- | --- | --- |
| TN | 0.142210795 | 0.140692476 |
| Her2 | 0.126142554 |  |

## Additional file 4: Giant California sea cucumber (*Parastichopus californicus*) example

The dataset used for assignment of the giant California sea cucumber comprised 717 individuals from the northeastern Pacific Ocean coast. The genotype of each individual was described by 3 699 SNPs [1]. The format of the data file used by the MMD method is described in Additional file 1: Supplementary data file S5.

In Ref. [1], a leave-one out strategy [2] was used for self-attribution of *P. californicus* to the north and south regions of the coast. The leave-one out strategy consists in removing an individual from the dataset which is attributed to the sources that are described by the remaining genotypes. As for the rest of examples studied in this work, we used a Monte-Carlo crossvalidation strategy [2] for self-attribution with the MMD method. In particular, we removed pairs of individuals (i.e. *I_u_* = 2) whose origin was assumed to be unknown. This procedure was repeated 100 times by randomly removing pairs of individuals from each region.

The probability of correct self-attribution *p*^sa^ for a removed pair was estimated for each realisation. Individuals from the north were correctly attributed with probability *p*^sa^ = 1 in most of the realisations (see the histogram for *p*^sa^ in Fig. AF4.1(a)). On average, the self-attribution accuracy for the north region was 92%. This is close to the 90% accuracy reported for the leave-one-out method in [1]. The probability of correct self-attribution for individuals from the south region is more widely spread than that for the north. It ranges between ~ 0.6 and 1 (Fig. AF4.1(b)) which is statistically compatible with the 88% correct self-attribution reported in Ref. [1]. The mean self-attribution accuracy in this case is 71%.

With regards to the selection of informative *P. californicus* loci, self-attribution is slightly more accurate with strategy S1 but differences are not significant for selections of more than 100 SNPs (Fig. AF4.2). An overall self-attribution accuracy of 76% was achieved by selecting the 100 most informative loci with strategy S1 (this accuracy is only ~ 8% lower than that obtained with all loci). This trend is statistically compatible with the findings in [1] which reported a self-attribution success of ~ 80% when selecting 100 SNPs.

---

**Fig. AF4.1.**
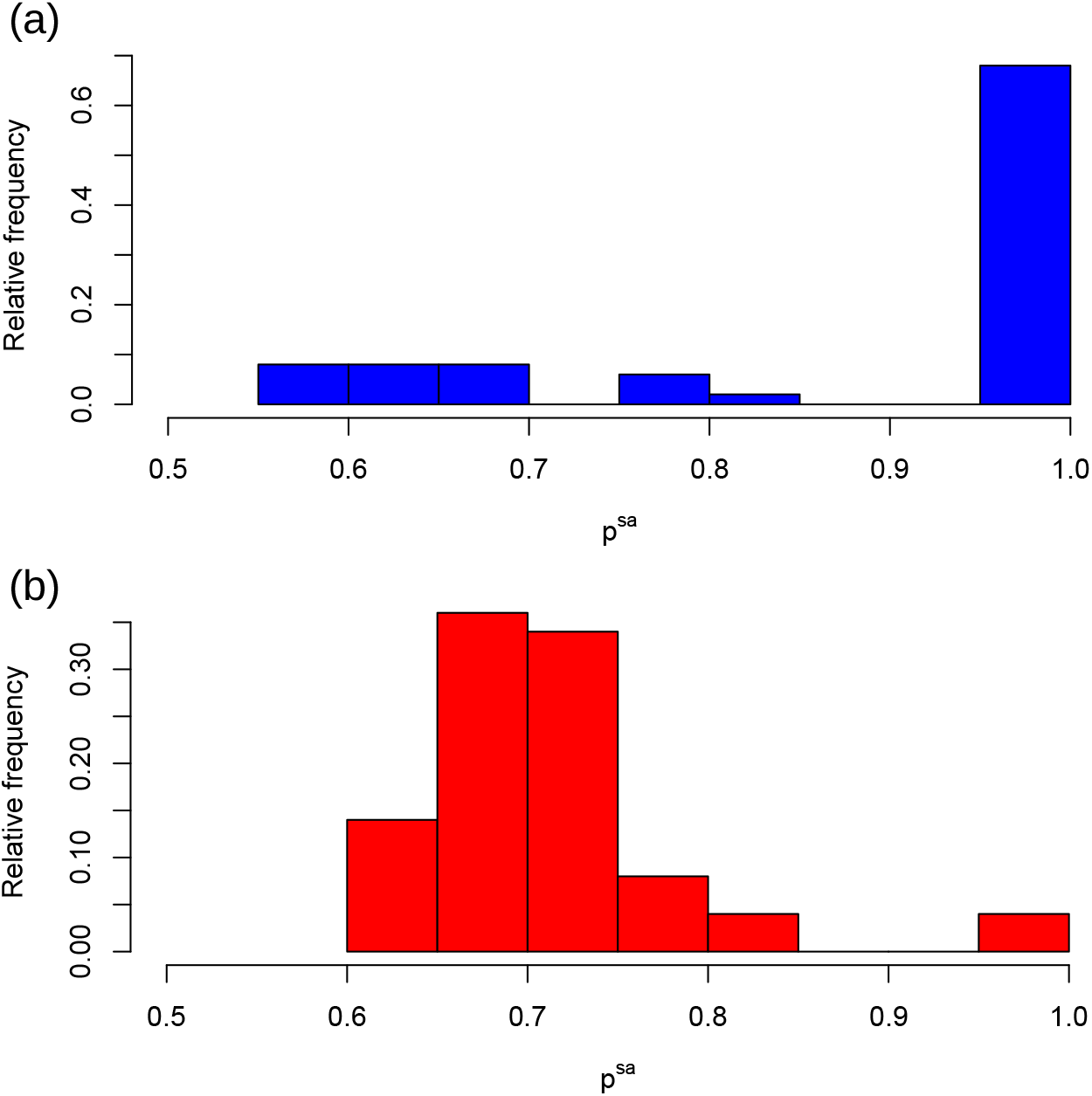
Correct self-attribution of *P. californicus*. Histograms show the relative frequency of the probability of correct self-attribution, *p*^sa^, of samples from (a) north and (b) south regions obtained by randomly removing pairs of individuals from each region.

**Fig. AF4.2.**
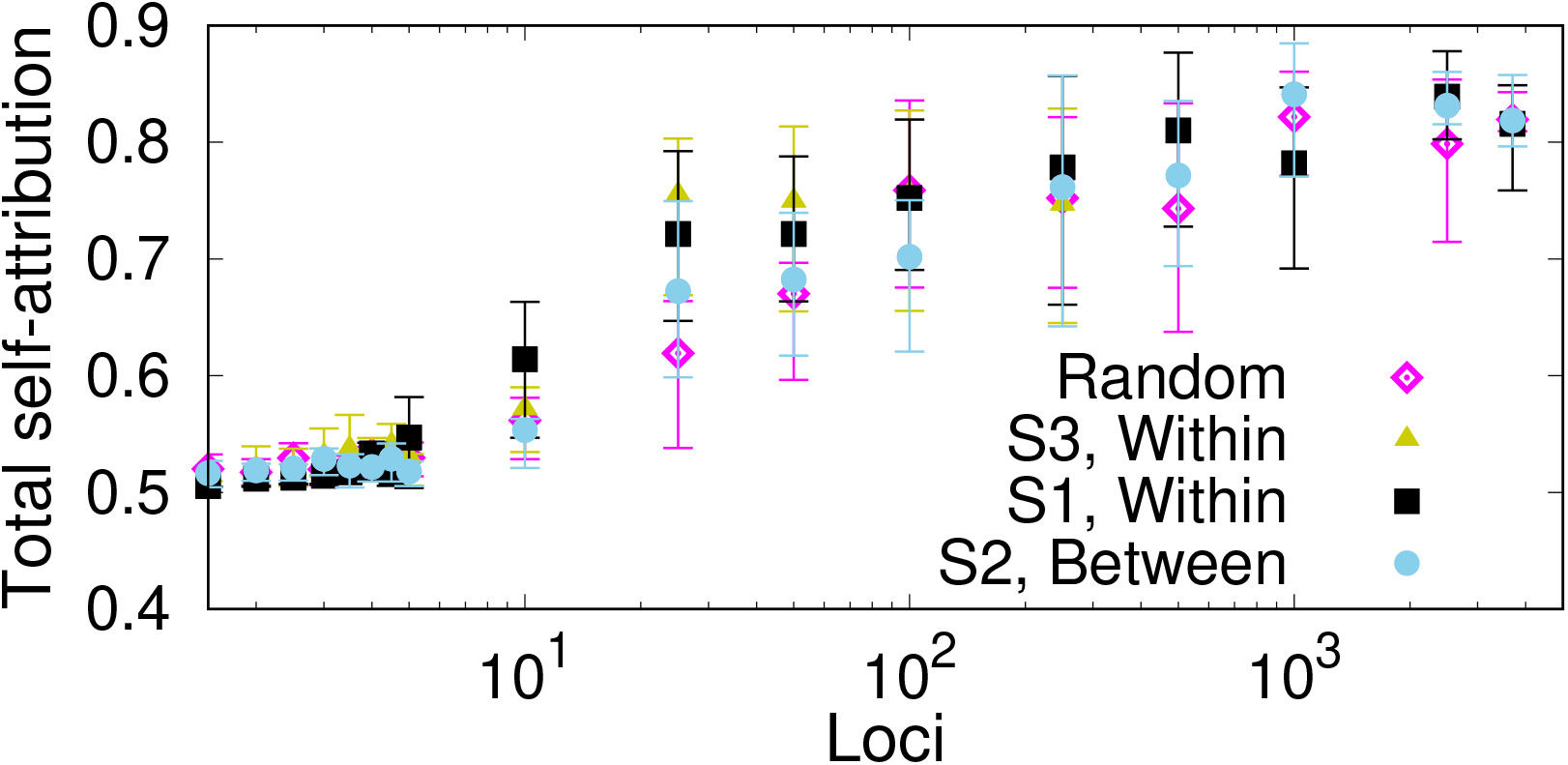
Selection of markers for self-attribution of *P. californicus* marine cucumber genotypes. Symbols show the probability of correct self-attribution, *p*^sa^, for samples from the north and south regions of the northeastern Pacific Ocean North American coast. The probability is plotted as a function of the number of SNPs selected at random and with strategies S1 (loci ranked in decreasing within-source diversity), S2 (loci ranked in decreasing between-source diversity) and S3 (reordering the loci ranking of S1 to reduce loci redundancy). Results were obtained by removing pairs of samples from sources to be attributed as if their origin was unknown. The process was repeated 100 times for each selected pair.

## Additional file 5: Breast cancer proteomic example

The dataset used in the breast cancer example comprised 40 breast cancer samples of three subtypes: 14 oestrogen receptor and/or progesterone receptor positive (ERPR positive) cases, 15 epidermal growth factor receptor *ErbB2/Her2* positive (Her2 positive) cases and 11 triple negative (TN) cases [1]. For each sample, the data from Ref. [1] provide the mass spectrum intensity *I_MS_* detected at 65 533 discrete values of *m/z* (ionic mass per unit charge). In order to represent these data as a feature vector suitable for MMD, the mass spectrum for each sample was transformed by replacing positive values of *I_MS_* by 1. This resulted in a feature vector of 65 533 elements with values 0 or 1 which defines a proteotype for each sample (Additional file 1: Suppl data file S6). The feature vector for each sample defines a multilocus proteotype analogous to the multilocus genotypes used in the *Campylobater*, human and *P. californicus* examples.

Self-attribution was performed by a Monte-Carlo cross-validation strategy [2] similar to that used for *P. californicus*, i.e. *I_u_* = 2 samples were randomly removed whose cancer subtypes were assumed to be unknown. This procedure is repeated for 100 different selections of pairs. Since the number of samples in the proteomic dataset is relatively small (40 samples), removing few samples is important to make sure that the remaining samples represent the sources as accurately as possible.

Overall, cancer samples were correctly attributed to their subtype (ERPR, Her2 or TN) in 63% of the cases. The average self-attribution probabilities for ERPR, Her2 and TN tumours were 0.64, 0.57 and 0.68, respectively (see Fig. AF5.1). Wrong self-attribution of any of the subtypes was approximately evenly distributed among the two wrong subtypes.

Self-attribution of breast cancer tumours is not significantly affected by the strategy used to select loci (Fig. AF5.2). Attribution accuracy saturates for selections of more than ~ 500 loci irrespective of the strategy used for loci selection.

---

**Fig. AF5.1.**
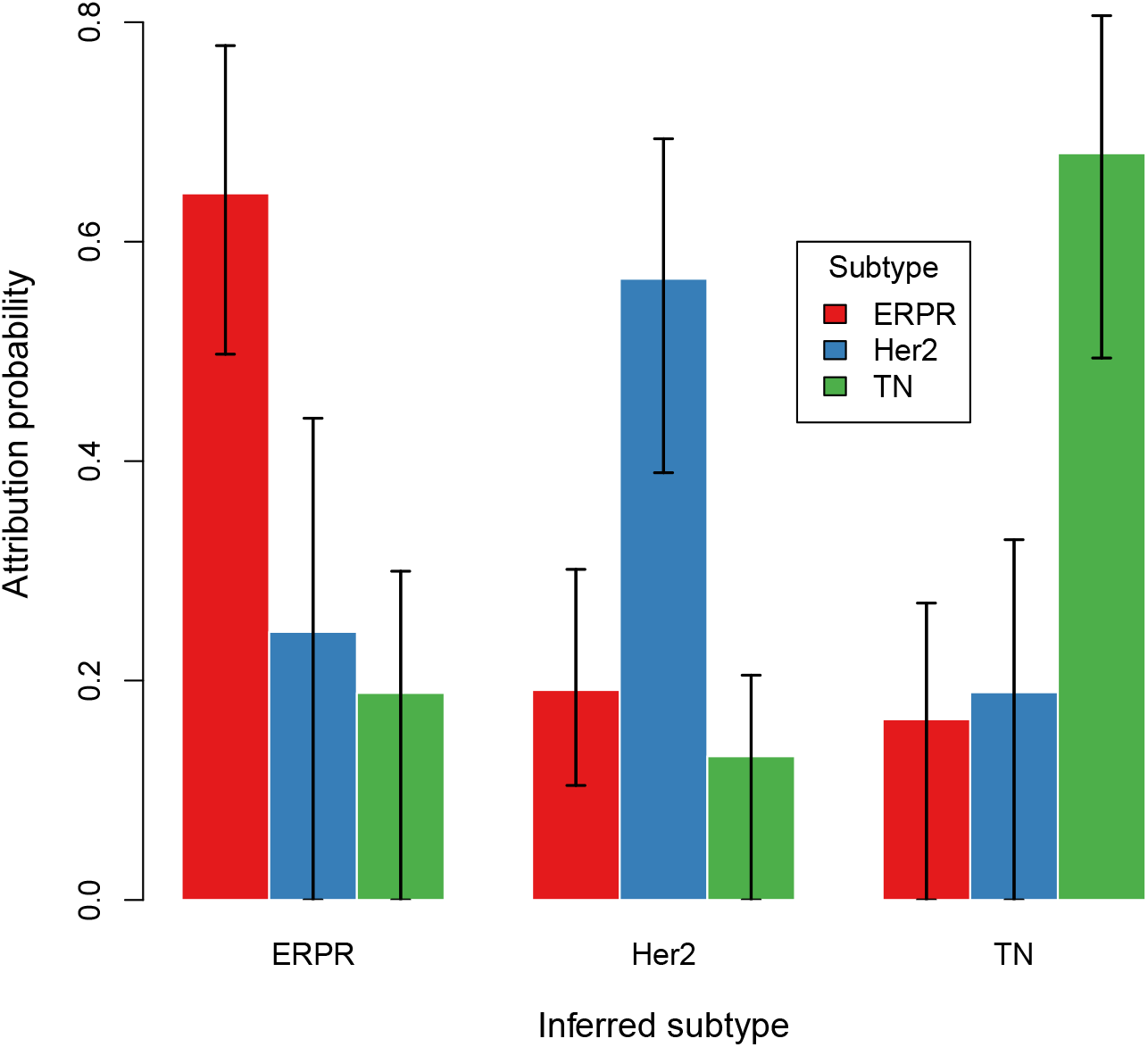
Self-attribution of breast cancer tumours based on proteomic data. Each sample (40 in total) is described by a 65 533 loci proteotype. Different colours, indicated in the legend, correspond to different cancer subtypes (ERPR, Her2 and TN). The bars for a given subtype provide the probability *p_u,s_* that removed samples, u, from this subtype are attributed to each of the possible sources, *s*. The probability indicated by the bars corresponds to the mean assignment probability over different selections. On average, ERPR, Her2 and TN subtypes are correctly attributed in 64%, 57% and 68% of the cases, respectively.

**Fig. AF5.2.**
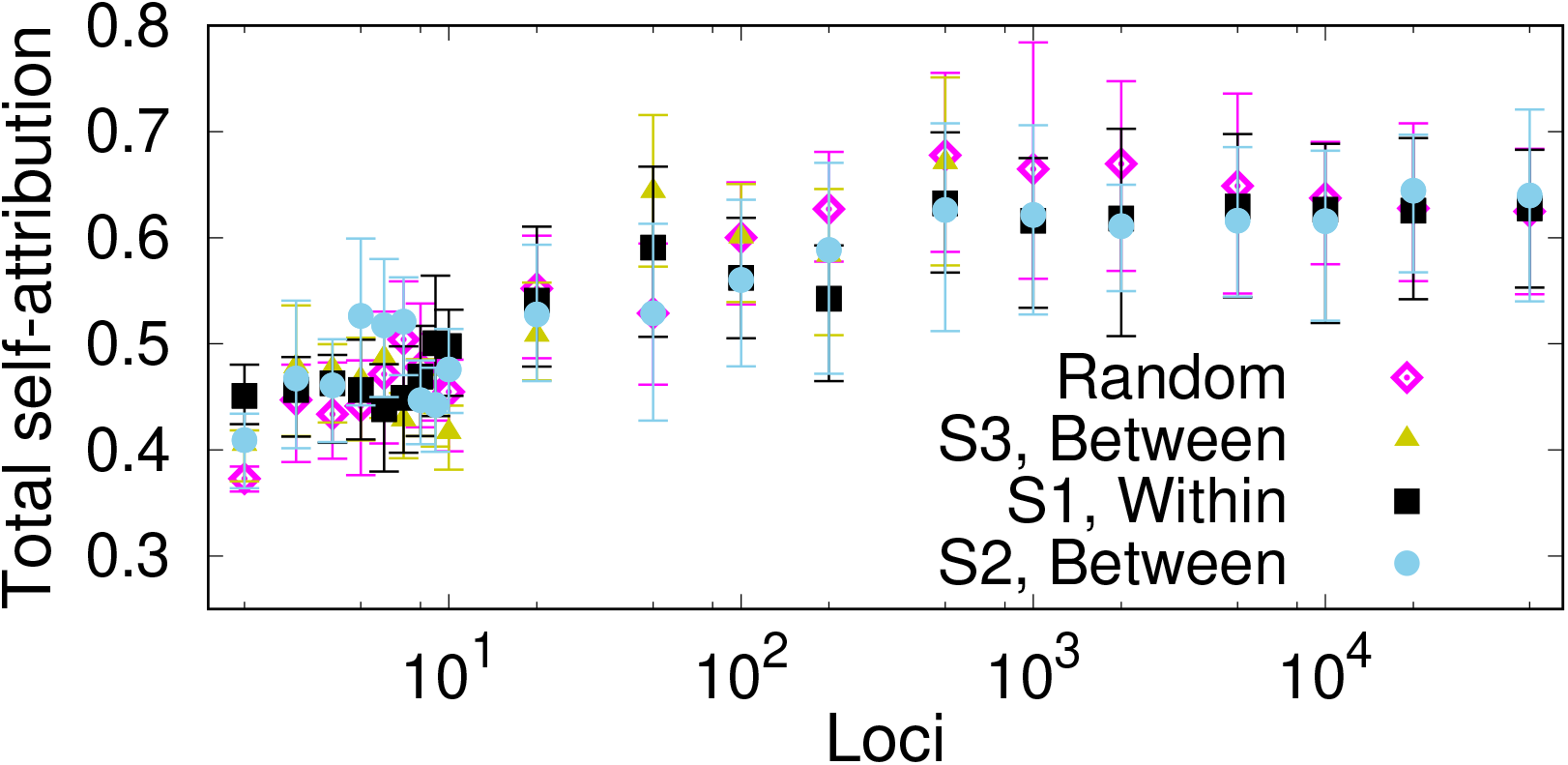
Selection of markers for self-attribution of breast cancer proteotypes. Symbols show the self-attribution probability *p*^sa^ that individuals from any of the three cancer subtypes (ERPR, Her2 or TN) are correctly attributed to their source. The probability is plotted as a function of the number of SNPs selected at random and with strategies S1 (loci ranked in decreasing within-source diversity), S2 (loci ranked in decreasing between-source diversity) and S3 (reordering the loci ranking of S2 to reduce loci redundancy).

## Additional file 6: Comparison of the MMD with other methods - Analytical considerations

### I. MMD METHOD IN TERMS OF ALLELE PROBABILITIES

Here, we present a description of the MMD method in terms of allele probabilities which is useful to compare with assignment methods that rely on allele probabilities. Our description applies to the particular case in which genotypes consist of *L unlinked* loci with two alleles each. Under this assumption, the Hamming distance *d*_H_(**u, a**_*i,s*_) is a random variable obeying a Poisson’s Binomial distribution [1] with success probabilities 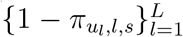. Here, *π_u_l_,l,s_* is the probability that the allele *u_l_* in the individual to be assigned is observed at locus *l* in source *s*. In general, *π_a,l,s_* denotes the probability of allele *a* at locus *l* in population *s*.

The measures of similarity between individuals and sources used in previous work based on allele frequencies can be viewed as *particular characteristics of the Hamming distance distribution* used by the MMD method. For instance, the likelihood function,

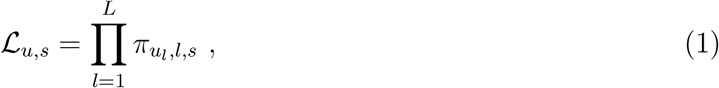

used in many assignment tests [2–7] corresponds to the probability that *d*_H_(**u, a**_*i,s*_) = 0, i.e. the probability that the genotype **u** exists in source *s*. Genetic distances used in distance-based assignment tests [4, 5], can also be expressed in terms of the probabilities {*π_u_l_,l,s_*}. For example, Nei’s *D_A_* distance [8] between the individual to be assigned and source *s* is

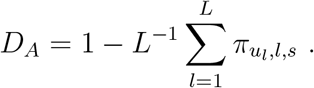

We note that some classical genetic distances [8] such as Nei’s standard genetic distance, *D_S_*, or Nei’s minimum genetic distance, *D_m_*, depend on the gene identity [9] of the sources, 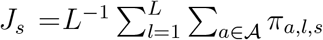, in addition to the probabilities {*π_u_l_,l,s_*}. For example, Nei’s standard genetic distance between **u** and source *s* is

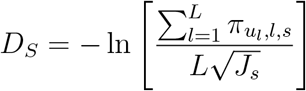

The gene identity is intrinsic to sources and does not reflect the similarity between the individual to be attributed and sources. In general, methods based on *D_S_* and *D_m_* will predict a higher attribution to the source with lower gene identity but this has nothing to do with the individual to be attributed.

### II. ATTRIBUTION ERRORS ASSOCIATED WITH ERRORS IN ALLELE PROBABILITIES

As mentioned in the main text, errors in the estimates of allele probabilities {*π_a,l,s_*} used to characterise sources will induce an error in attribution. Here we estimate the dependence of the attribution error on the number *L* of loci in the genotypes and the number *I_s_* of genotypes used to describe each source.

#### A. Attribution error for the MMD method

For the MMD method, errors in the estimates of the allele probabilities propagate to the quantile λ_*u,s*_(*q*), score *σ_u,s_* and attribution probability *p_u,s_* defined in the Methods of the main text. The dependence of the errors of λ_*u,s*_(*q*) and *σ_u,s_* on *L* and *I_s_* can be estimated for a simple model for unlinked loci in which alleles have the same probability distribution for all loci, i.e. a model with *π_u_l_,l,s_* = *r_s_* independently of *l*. In this case, the Hamming distance obeys a binomial distribution for *L* Bernoulli trials with probability of success 1 – *r_s_*. In the limit of large *L*, the binomial distribution can be approximated by a normal distribution with mean *μ_s_* = *L*(1 – *r_s_*) and variance 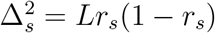. Under these assumptions, the quantile λ_*u,s*_(*q*) satisfies

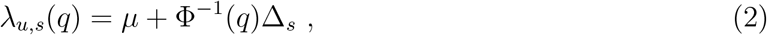

and the score *σ_u,s_* quantifying the proximity of genotype **u** to source *s* is

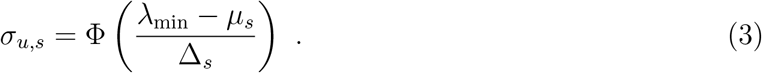

Here, λ_min_ = min_*s*_{λ_*u,s*_(*q*)} and Φ^−1^(*x*) is the inverse of the cumulative distribution function for the standard normal distribution.

From Eq. (2), the error of λ_*u,s*_(*q*) in the limit of extended genotypes with large *L* is given by

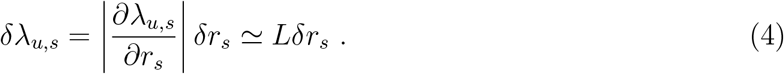

Here, *δr_s_* is the error in the allele probabilities. In the MMD method and other methods that approximate these probabilities by the observed allele frequencies, the error is 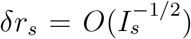. Therefore,

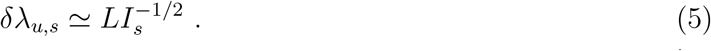

Since λ_*u,s*_ ≃ *L* (cf. Eq. (2)), we conclude that the relative error of λ_*u,s*_ is 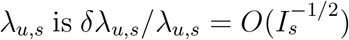, i.e. it does not increase with the number of loci, *L*.

Let us denote the closest source to individual **u** as *s*_closest_ (this is the source with λ_*u,s*_closest__ = λ_min_). From Eq. (3), the error in the assignment score *σ_u,s_* is given by:

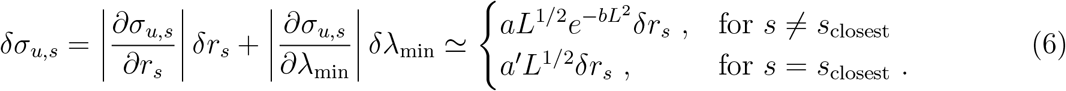

Here, *a, a′* and *b* are independent of *L* and we have assumed that *δr_s_* is approximately the same for all sources, including *s*_closest_. One can show that the error for the attribution probability *p_u,s_* is proportional to that of *δσ_u,s_*.

To summarise, our arguments show that the assignment error for the MMD method is *O*(*L*^1/2^). In the particular case in which the allele probabilities are estimated by frequencies, one has 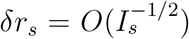 and the assignment error is 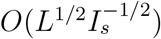, i.e. it increases with *L* and decreases with the number of genotypes used to define the sources.

#### B. Attribution error for the likelihood function

The error for the likelihood function given by Eq. (1) can be easily calculated as a function of the errors {*δπ_u_l_;l;s_*} for the allele probabilities. Propagation of errors gives

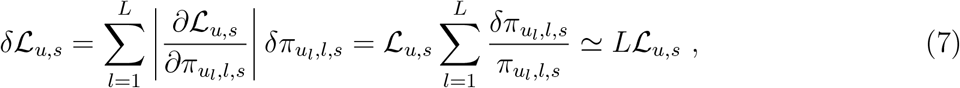

where we have assumed *δπ_u_l_,l,s_* > 0 for all loci.

According to Eq. (7), the relative error of the likelihood function, 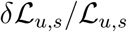, increases with *L* unless the errors in the probability estimates, {*δπ_u_l_,l,s_*}, are zero.

The log-likelihood function is more commonly used than the likelihood itself. One can easily show that the error for the log-likelihood function typically equals the relative error of 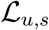 and is therefore *O*(*L*). This shows that attribution errors based on a likelihood function increase faster with *L* than those for the MMD method.

## Additional file 7: Exclusion method

Here, we apply the threshold exclusion method proposed in [1] to the MMD source attribution results. The exclusion method consists in setting a threshold *T* for the probability *p_u,s_* such that an individual *u* is assigned to source *s* if *p_u,s_ ≥ T*. Otherwise, if *p_u,s_ < T*, the individual cannot be attributed to the source *s*. When *p_u,s_ < T* for all the sampled sources, 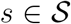, the individual is not assigned to any source. The performance of this method was explored to test the MMD attribution results for human genotypes with 659 276 SNPs and *Campylobacter* genotypes with 25 937 SNPs. For the human example, we focused on self-attribution to 7 geographical regions. In this case, a very small proportion of genotypes are excluded even for very selective values of *T* (see Fig. AF7.1). In fact, only 2% of individuals are excluded from all regions for T = 1. In fact, for T =1, all individuals are attributed to the correct source except for 10% of individuals that are excluded from Middle East and 1% that are excluded from C/S Asia. To some extent, the high accuracy found for this dataset could be expected since we are dealing with self-attribution and the true region of individuals has been sampled for sure.

Exclusion is more prominent for *Campylobacter* isolates (see Fig. AF7.2). For *T* > 0.9, more than 60% of isolates from human patients are excluded from all sources. Since the origin of human isolates is unknown, one could conclude that there is a high percentage of isolates that originated from sources that were not sampled. However, this is not a solid conclusion since the method also predicts high exclusion percentages (> 37% on average for *T* > 0.9) for isolates from food and animal sources whose true source is known. The exclusion percentage is particularly high for sheep (all isolates excluded from all sources for *T* > 0.7) and cattle (52% excluded for any *T* > 0.8). High exclusion rates in this example are likely due to a low genetic differentiation between sources. In this situation, forcing assignment to a single source is not well justified.

---

**Fig. AF7.1.**
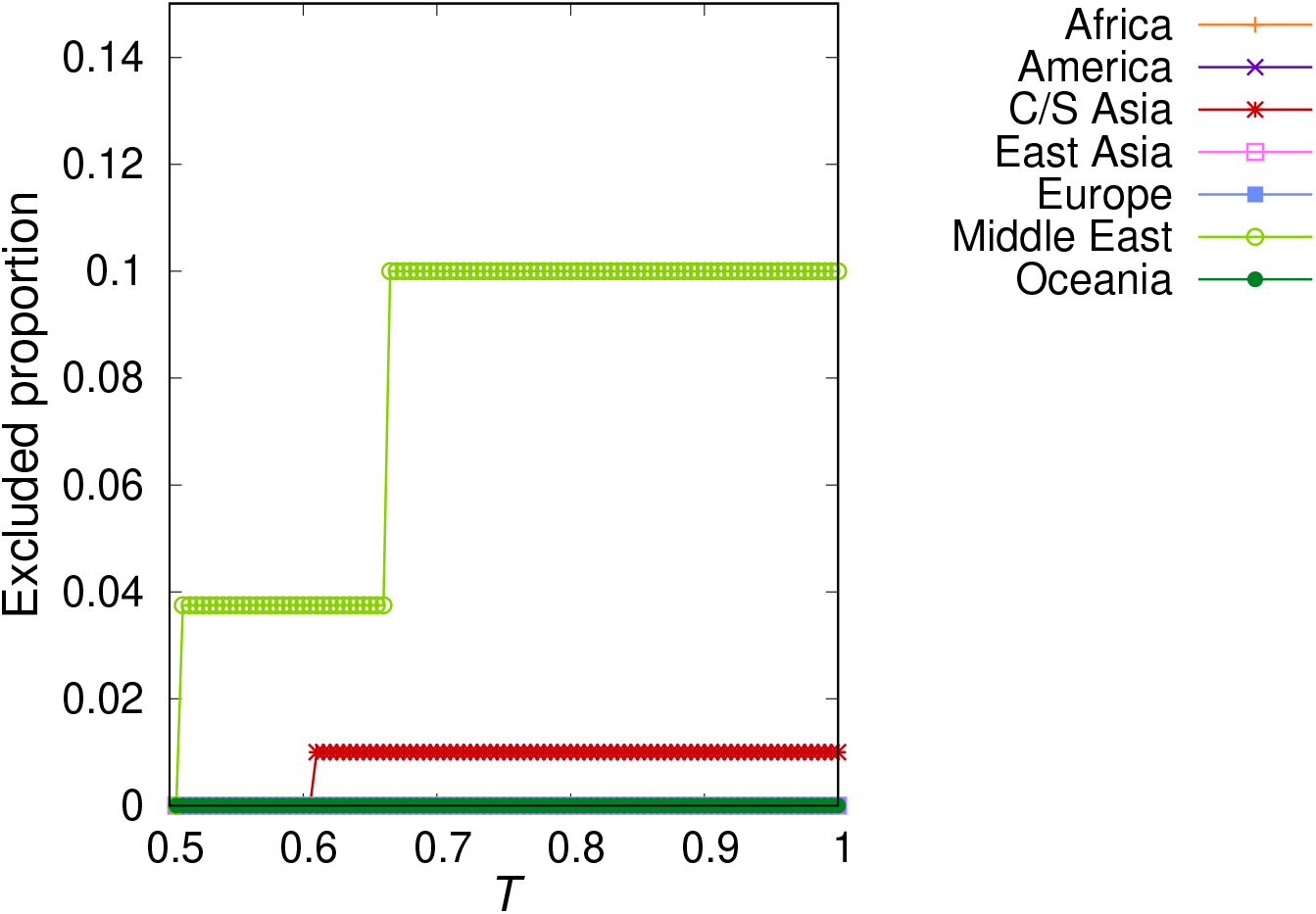
Exclusion test for humans based on 659 276 SNP genotypes. For a given geographical region, the proportion of individuals that are not attributed to any region (i.e. individuals with *p_u,s_ < T* for all regions, s) is plotted as a function of the exclusion threshold, *T*. Different symbols correspond to individuals from different regions, as marked by the legend.

**Fig. AF7.2.**
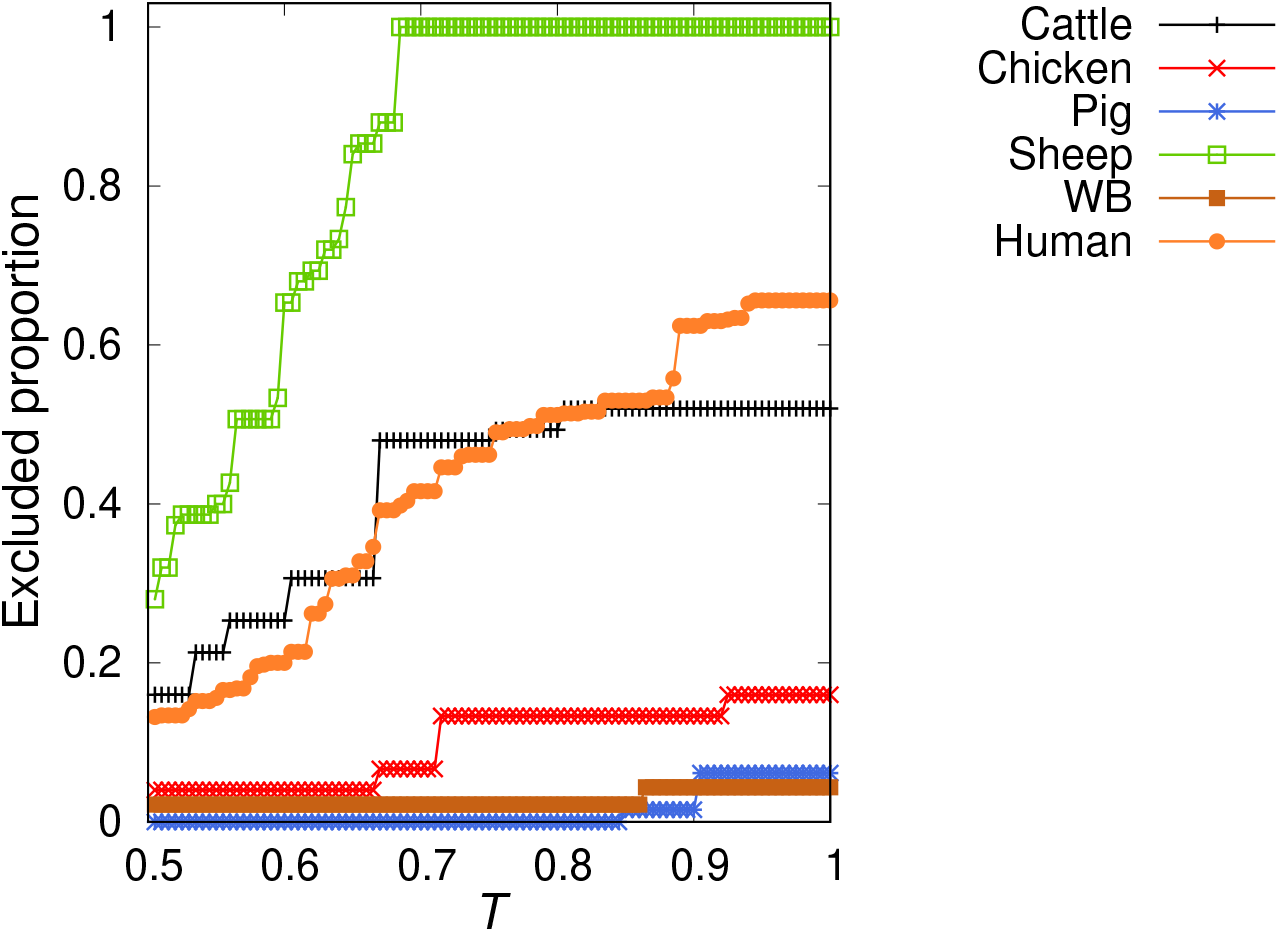
Exclusion test for Campylobacter isolates based on 25938 _cg_SNP genotypes. For a given Campylobacter reservoir, the proportion of isolates that are not attributed to any source (i.e. isolates with *p_u,s_ < T* for all sources) is plotted as a function of the exclusion threshold, *T*. Different symbols correspond to isolates from different reservoirs, as marked by the legend.

## References

1. Waser, P.M., Strobeck, C.: Genetic signatures of interpopulation dispersal. Trends in ecology & evolution 13(2), 43–4 (1998). doi: 10.1016/S0169-5347(97)01255-X

2. Davies, N., Villablanca, F.X., Roderick, G.K., Davies, N., Villablanca, F.X., Roderick, G.K., Davies, N., Villablanca, F.X., Roderick, G.K.: Determining the source of individuals: multilocus genotyping in nonequilibrium population genetics. Trends in ecology & evolution 14(1), 17–21 (1999). doi:10.1016/S0169-5347(98)01530-4

3. Paetkau, D., Calvert, W., Stirling, I., Strobeck, C.: Microsatellite analysis of population structure in Canadian polar bears. Molecular Ecology 4(3), 347–354 (1995). doi:10.1111/j.1365-294X.1995.tb00227.x

4. Nielsen, R., Mattila, D.K., Clapham, P.J., Palsbøll, P.J.: Statistical approaches to paternity analysis in natural populations and applications to the North Atlantic humpback whale. Genetics 157(4), 1673–1682 (2001)

5. Manel, S., Berthier, P., Luikart, G.: Detecting Wildlife Poaching: Identifying the Origin of Individuals with Bayesian Assignment Tests and Multilocus Genotypes. Conservation Biology 16(3), 650–659 (2002). doi:10.1046/j.1523-1739.2002.00576.x

6. Berry, O., Tocher, M.D., Sarre, S.D.: Can assignment tests measure dispersal? Molecular Ecology 13(3), 551–561 (2004). doi:10.1046/j.1365-294X.2004.2081.x

7. Storer, C.G., Pascal, C.E., Roberts, S.B., Templin, W.D., Seeb, L.W., Seeb, J.E.: Rank and Order: Evaluating the Performance of SNPs for Individual Assignment in a Non-Model Organism. PLoS ONE 7(11), 49018 (2012). doi:10.1371/journal.pone.0049018

8. McCarthy, N.D., Colles, F.M., Dingle, K.E., Bagnall, M.C., Manning, G., Maiden, M.C.J., Falush, D.: Host-associated genetic import in Campylobacter jejuni. Emerging infectious diseases 13(2), 267–72 (2007). doi:10.3201/eid1302.060620

9. Pires, S.M., Evers, E.G., van Pelt, W., Ayers, T., Scallan, E., Angulo, F.J., Havelaar, A., Hald, T.: Attributing the human disease burden of foodborne infections to specific sources. Foodborne Pathogens and Disease 6(4), 417–424 (2009). doi:10.1089/fpd.2008.0208

10. EFSA: Scientific Opinion on the evaluation of molecular typing methods for major food-borne microbiological hazards and their use for attribution modelling, outbreak investigation and scanning surveillance: Part 1 (evaluation of methods and applications). EFSA Journal 11(12), 3502 (2013). doi:10.2903/j.efsa.2013.3502

11. Sheppard, S.K., Dallas, J.F., Strachan, N.J.C., MacRae, M., McCarthy, N.D., Wilson, D.J., Gormley, F.J., Falush, D., Ogden, I.D., Maiden, M.C.J., Forbes, K.J.: Campylobacter genotyping to determine the source of human infection. Clinical infectious diseases: an official publication of the Infectious Diseases Society of America 48(8), 1072–8 (2009). doi: 10.1086/597402

12. Strachan, N.J.C., Gormley, F.J., Rotariu, O., Ogden, I.D., Miller, G., Dunn, G.M., Sheppard, S.K., Dallas, J.F., Reid, T.M.S., Howie, H., Maiden, M.C.J., Forbes, K.J.: Attribution of Campylobacter infections in northeast Scotland to specific sources by use of multilocus sequence typing. The Journal of infectious diseases 199(8), 1205–8 (2009). doi:10.1086/597417

13. Kittl, S., Heckel, G., Korczak, B.M., Kuhnert, P.: Source attribution of human Campylobacter isolates by MLST and Fla-typing and association of genotypes with quinolone resistance. PLoS ONE 8(11), 81796 (2013). doi:10.1371/journal.pone.0081796

14. Wilson, D.J., Gabriel, E., Leatherbarrow, A.J.H., Cheesbrough, J., Gee, S., Bolton, E., Fox, A., Fearnhead, P., Hart, C.A., Diggle, P.J.: Tracing the Source of Campylobacteriosis. PLoS Genetics 4(9), 1000203 (2008). doi:10.1371/journal.pgen.1000203

15. Mullner, P., Jones, G., Noble, A., Spencer, S.E.F., Hathaway, S., French, N.P.: Source attribution of food-borne zoonoses in New Zealand: A modified hald model. Risk Analysis 29(7), 970–984 (2009). doi:10.1111/j.1539-6924.2009.01224.x

16. Mughini Gras, L., Smid, J.H., Wagenaar, J.A., de Boer, A.G., Havelaar, A.H., Friesema, I.H.M., French, N.P., Busani, L., van Pelt, W.: Risk Factors for Campylobacteriosis of Chicken, Ruminant, and Environmental Origin: A Combined Case-Control and Source Attribution Analysis. PLoS ONE 7(8), 42599 (2012). doi:10.1371/journal.pone.0042599

17. Boysen, L., Rosenquist, H., Larsson, J.T., Nielsen, E.M., Sorensen, G., Nordentoft, S., Hald, T.: Source attribution of human campylobacteriosis in Denmark. Epidemiology and Infection 142(8), 1599–1608 (2014). doi: 10.1017/S0950268813002719

18. Rosner, B.M., Schielke, A., Didelot, X., Kops, F., Breidenbach, J., Willrich, N., Gölz, G., Alter, T., Stingl, K., Josenhans, C., Suerbaum, S., Stark, K.: A combined case-control and molecular source attribution study of human Campylobacter infections in Germany, 2011-2014. Scientific Reports 7(1), 5139 (2017). doi:10.1038/s41598-017-05227-x

19. Miller, P., Marshall, J., French, N., Jewell, C.: sourceR: Classification and source attribution of infectious agents among heterogeneous populations. PLOS Computational Biology 13(5), 1005564 (2017). doi:10.1371/journal.pcbi.1005564

20. Rosenberg, N.A., Pritchard, J.K., Weber, J.L., Cann, H.M., Kidd, K.K., Zhivotovsky, L.A., Feldman, M.W.: Genetic structure of human populations. Science 298(5602), 2381–2385 (2002). doi:10.1126/science.1078311

21. Rosenberg, N.A., Li, L.M., Ward, R., Pritchard, J.K.: Informativeness of Genetic Markers for Inference of Ancestry. The American Journal of Human Genetics 73(6), 1402–1422 (2003). doi: 10.1086/380416

22. Pemberton, T.J., DeGiorgio, M., Rosenberg, N.A.: Population Structure in a Comprehensive Genomic Data Set on Human Microsatellite Variation. G3 Genes|Genomes|Genetics 3(5), 891–907 (2013). doi:10.1534/g3.113.005728

23. Faria, D.A., Mamani, E.M.C., Pappas, G.J., Grattapaglia, D.: Genotyping systems for Eucalyptus based on tetra-, penta-, and hexanucleotide repeat EST microsatellites and their use for individual fingerprinting and assignment tests. Tree Genetics & Genomes 7(1), 63–77 (2011). doi:10.1007/s11295-010-0315-9

24. Maiden, M.C.J., van Rensburg, M.J.J., Bray, J.E., Earle, S.G., Ford, S.A., Jolley, K.A., McCarthy, N.D.: MLST revisited: the gene-by-gene approach to bacterial genomics. Nature Reviews Microbiology 11(10), 728–736 (2013). doi: 10.1038/nrmicro3093

25. Nielsen, E.M., Bjorkman, J.T., Kiil, K., Grant, K., Dallman, T., Painset, A., Amar, C., Roussel, S., Guillier, L., Félix, B., Rotariu, O., Perez-Reche, F., Forbes, K., Strachan, N.: Closing gaps for performing a risk assessment on Listeria monocytogenes in ready to eat (RTE) foods: activity 3, the comparison of isolates from different compartments along the food chain, and from humans using whole genome sequencing (WGS) analysis. EFSA Supporting Publications 14(2) (2017). doi: 10.2903/sp.efsa.2017.EN-1151

26. Massung, R.F., Liu, L.-I., Qi, J., Knight, J.C., Yuran, T.E., Kerlavage, A.R., Parsons, J.M., Venter, J.C., Esposito, J.J.: Analysis of the Complete Genome of Smallpox Variola Major Virus Strain Bangladesh-1975. Virology 201(2), 215–240 (1994). doi: 10.1006/VIRO.1994.1288

27. Fouts, D.E., Mongodin, E.F., Mandrell, R.E., Miller, W.G., Rasko, D.A., Ravel, J., Brinkac, L.M., DeBoy, R.T., Parker, C. T., Daugherty, S.C., Dodson, R.J., Durkin, A.S., Madupu, R., Sullivan, S.A., Shetty, J.U., Ayodeji, M.A., Shvartsbeyn, A., Schatz, M.C., Badger, J.H., Fraser, C.M., Nelson, K.E.: Major Structural Differences and Novel Potential Virulence Mechanisms from the Genomes of Multiple Campylobacter Species. PLoS Biology 3(1), 15 (2005). doi: 10.1371/journal.pbio.0030015

28. Adams, M.D., Celniker, S.E., Holt, R.A., Evans, C.A., Gocayne, J.D., Amanatides, P.G., Scherer, S.E., Li, P.W., Hoskins, R.A., Galle, R.F., George, R.A., Lewis, S.E., Richards, S., Ashburner, M., Henderson, S.N., Sutton, G.G., Wortman, J.R., Yandell, M.D., Zhang, Q., Chen, L.X., Brandon, R.C., Rogers, Y.-H.C., Blazej, R.G., Champe, M., Pfeiffer, B.D., Wan, K.H., Doyle, C., Baxter, E.G., Helt, G., Nelson, C.R., Gabor, G.L., Miklos, Abril, J.F., Agbayani, A., An, H.-J., Andrews-Pfannkoch, C., Baldwin, D., Ballew, R.M., Basu, A., Baxendale, J., Bayraktaroglu, L., Beasley, E.M., Beeson, K.Y., Benos, P.V., Berman, B.P., Bhandari, D., Bolshakov, S., Borkova, D., Botchan, M.R., Bouck, J., Brokstein, P., Brottier, P., Burtis, K.C., Busam, D.A., Butler, H., Cadieu, E., Center, A., Chandra, I., Cherry, J.M., Cawley, S., Dahlke, C., Davenport, L.B., Davies, P., de Pablos, B., Delcher, A., Deng, Z., Mays, A.D., Dew, I., Dietz, S.M., Dodson, K., Doup, L.E., Downes, M., Dugan-Rocha, S., Dunkov, B.C., Dunn, P., Durbin, K.J., Evangelista, C.C., Ferraz, C., Ferriera, S., Fleischmann, W., Fosler, C., Gabrielian, A.E., Garg, N.S., Gelbart, W.M., Glasser, K., Glodek, A., Gong, F., Gorrell, J.H., Gu, Z., Guan, P., Harris, M., Harris, N.L., Harvey, D., Heiman, T.J., Hernandez, J.R., Houck, J., Hostin, D., Houston, K.A., Howland, T.J., Wei, M.-H., Ibegwam, C., Jalali, M., Kalush, F., Karpen, G.H., Ke, Z., Kennison, J.A., Ketchum, K.A., Kimmel, B.E., Kodira, C.D., Kraft, C., Kravitz, S., Kulp, D., Lai, Z., Lasko, P., Lei, Y., Levitsky, A.A., Li, J., Li, Z., Liang, Y., Lin, X., Liu, X., Mattei, B., McIntosh, T.C., McLeod, M.P., McPherson, D., Merkulov, G., Milshina, N.V., Mobarry, C., Morris, J., Moshrefi, A., Mount, S.M., Moy, M., Murphy, B., Murphy, L., Muzny, D.M., Nelson, D.L., Nelson, D.R., Nelson, K.A., Nixon, K., Nusskern, D.R., Pacleb, J.M., Palazzolo, M., Pittman, G.S., Pan, S., Pollard, J., Puri, V., Reese, M.G., Reinert, K., Remington, K., Saunders, R.D.C., Scheeler, F., Shen, H., Shue, B.C., Sidén-Kiamos, I., Simpson, M., Skupski, M.P., Smith, T., Spier, E., Spradling, A.C., Stapleton, M., Strong, R., Sun, E., Svirskas, R., Tector, C., Turner, R., Venter, E., Wang, A.H., Wang, X., Wang, Z.-Y., Wassarman, D.A., Weinstock, G.M., Weissenbach, J., Williams, S.M., Woodage, T., Worley, K.C., Wu, D., Yang, S., Yao, Q.A., Ye, J., Yeh, R.-F., Zaveri, J.S., Zhan, M., Zhang, G., Zhao, Q., Zheng, L., Zheng, X.H., Zhong, F.N., Zhong, W., Zhou, X., Zhu, S., Zhu, X., Smith, H.O., Gibbs, R.A., Myers, E.W., Rubin, G.M., Venter, J.C.: The Genome Sequence of Drosophila melanogaster. Science 287(5461), 2185–2195 (2000)

29. Galagan, J.E., Calvo, S.E., Borkovich, K.A., Selker, E.U., Read, N.D., Jaffe, D., FitzHugh, W., Ma, L.-J., Smirnov, S., Purcell, S., Rehman, B., Elkins, T., Engels, R., Wang, S., Nielsen, C.B., Butler, J., Endrizzi, M., Qui, D., Ianakiev, P., Bell-Pedersen, D., Nelson, M.A., Werner-Washburne, M., Selitrennikoff, C.P., Kinsey, J.A., Braun, E.L., Zelter, A., Schulte, U., Kothe, G.O., Jedd, G., Mewes, W., Staben, C., Marcotte, E., Greenberg, D., Roy, A., Foley, K., Naylor, J., Stange-Thomann, N., Barrett, R., Gnerre, S., Kamal, M., Kamvysselis, M., Mauceli, E., Bielke, C., Rudd, S., Frishman, D., Krystofova, S., Rasmussen, C., Metzenberg, R.L., Perkins, D.D., Kroken, S., Cogoni, C., Macino, G., Catcheside, D., Li, W., Pratt, R.J., Osmani, S.A., DeSouza, C.P.C., Glass, L., Orbach, M.J., Berglund, J.A., Voelker, R., Yarden, O., Plamann, M., Seiler, S., Dunlap, J., Radford, A., Aramayo, R., Natvig, D.O., Alex, L.A., Mannhaupt, G., Ebbole, D.J., Freitag, M., Paulsen, I., Sachs, M.S., Lander, E.S., Nusbaum, C., Birren, B.: The genome sequence of the filamentous fungus Neurospora crassa. Nature 422(6934), 859–868 (2003). doi:10.1038/nature01554

30. Human Genome Sequencing Consortium, I.: Finishing the euchromatic sequence of the human genome. Nature 431(7011), 931–945 (2004). doi:10.1038/nature03001

31. Kao, R.R., Haydon, D.T., Lycett, S.J., Murcia, P.R.: Supersize me: How whole-genome sequencing and big data are transforming epidemiology. Trends in Microbiology 22(5), 282–291 (2014). doi: 10.1016/j.tim.2014.02.011

32. Bergholz, T.M., Moreno Switt, A.I., Wiedmann, M.: Omics approaches in food safety: Fulfilling the promise? Trends in Microbiology 22(5), 275–281 (2014). doi:10.1016/j.tim.2014.01.006

33. Harris, S.R., Feil, E.J., Holden, M.T.G., Quail, M.A., Nickerson, E.K., Chantratita, N., Gardete, S., Tavares, A., Day, N., Lindsay, J.A., Edgeworth, J.D., de Lencastre, H., Parkhill, J., Peacock, S.J., Bentley, S.D.: Evolution of MRSA During Hospital Transmission and Intercontinental Spread. Science 327(5964), 469–474 (2010)

34. Franz, E., Delaquis, P., Morabito, S., Beutin, L., Gobius, K., Rasko, D.A., Bono, J., French, N., Osek, J., Lindstedt, B.A., Muniesa, M., Manning, S., LeJeune, J., Callaway, T., Beatson, S., Eppinger, M., Dallman, T., Forbes, K.J., Aarts, H., Pearl, D. L., Gannon, V.P.J., Laing, C.R., Strachan, N.J.C.: Exploiting the explosion of information associated with whole genome sequencing to tackle Shiga toxin-producing Escherichia coli (STEC) in global food production systems. International Journal of Food Microbiology 187, 57–72 (2014). doi:10.1016/j.ijfoodmicro.2014.07.002

35. Strachan, N.J.C., Rotariu, O., Lopes, B., MacRae, M., Fairley, S., Laing, C., Gannon, V., Allison, L.J., Hanson, M.F., Dallman, T., Ashton, P., Franz, E., van Hoek, A.H.A.M., French, N.P., George, T., Biggs, P.J., Forbes, K.J.: Whole Genome Sequencing demonstrates that Geographic Variation of Escherichia coli O157 Genotypes Dominates Host Association. Scientific Reports 5, 14145 (2015). doi:10.1038/srep14145

36. Mughini-Gras, L., Kooh, P., Augustin, J.-C., David, J., Fravalo, P., Guillier, L., Jourdan-Da-Silva, N., Thébault, A., Sanaa, M., Watier, L., Diseases, T.A.W.G.o.S.A.o.F.: Source Attribution of Foodborne Diseases: Potentialities, Hurdles, and Future Expectations. Frontiers in Microbiology 9, 1983 (2018). doi:10.3389/fmicb.2018.01983

37. Pritchard, J.K., Stephens, M.M., Donnelly, P.: Inference of population structure using multilocus genotype data. Genetics 155(2), 945–959 (2000). doi:10.1111/j.1471-8286.2007.01758.x

38. Falush, D., Stephens, M., Pritchard, J.K.: Inference of population structure using multilocus genotype data: linked loci and correlated allele frequencies. Genetics 164, 1567–1587 (2003). doi:10.1111/j.1471-8286.2007.01758.x.1508.00973

39. Tang, H., Peng, J., Wang, P., Risch, N.J.: Estimation of individual admixture: Analytical and study design considerations. Genetic Epidemiology 28(4), 289–301 (2005). doi:10.1002/gepi.20064

40. Alexander, D.H., Novembre, J., Lange, K.: Fast model-based estimation of ancestry in unrelated individuals. Genome Research (2009). doi:10.1101/gr.094052.109

41. Raj, A., Stephens, M., Pritchard, J.K.: fastSTRUCTURE: Variational Inference of Population Structure in Large SNP Data Sets. Genetics 197(2), 573–589 (2014). doi:10.1534/genetics.114.164350

42. Lawson, D.J., Hellenthal, G., Myers, S., Falush, D.: Inference of Population Structure using Dense Haplotype Data. PLoS Genetics 8(1), 1002453 (2012). doi:10.1371/journal.pgen.1002453

43. Frichot, E., Mathieu, F., Trouillon, T., Bouchard, G., François, O.: Fast and efficient estimation of individual ancestry coefficients. Genetics 196(4), 973–83 (2014). doi:10.1534/genetics.113.160572

44. Beugin, M.P., Gayet, T., Pontier, D., Devillard, S., Jombart, T.: A fast likelihood solution to the genetic clustering problem. Methods in Ecology and Evolution 9(4), 1006–1016 (2018). doi:10.1111/2041-210X.12968.0608246v3

45. Patterson, N., Price, A.L., Reich, D.: Population Structure and Eigenanalysis. PLoS Genetics 2(12), 190 (2006). doi:10.1371/journal.pgen.0020190

46. Jombart, T., Devillard, S., Balloux, F.: Discriminant analysis of principal components: a new method for the analysis of genetically structured populations. BMC Genetics 11(1), 94 (2010). doi: 10.1186/1471-2156-11-94

47. Murphy, K.P.: Machine Learning: A Probabilistic Perspective. MIT Press, Cambridge, MA (2012)

48. Alexander, D.H., Lange, K.: Enhancements to the ADMIXTURE algorithm for individual ancestry estimation. BMC Bioinformatics 12(1), 246 (2011). doi:10.1186/1471-2105-12-246

49. Hellenthal, G., Busby, G.B.J., Band, G., Wilson, J.F., Capelli, C., Falush, D., Myers, S.: A genetic atlas of human admixture history. Science 343(6172), 747 (2014). doi:10.1126/science.1243518

50. Smouse, P.E., Chevillon, C.: Analytical Aspects of Population-Specific DNA Fingerprinting for Individuals. Journal of Heredity 89, 143–150 (1998)

51. Cornuet, J.M., Piry, S., Luikart, G., Estoup, A., Solignac, M.: New methods employing multilocus genotypes to select or exclude populations as origins of individuals. Genetics 153(4), 1989–2000 (1999). doi: 10.1038/368455a0

52. Rosenberg, N.A., Burke, T., Elo, K., Feldman, M.W., Freidlin, P.J., Groenen, M.A.M., Hillel, J., Mäki-Tanila, A., Tixier-Boichard, M., Vignal, A., Wimmers, K., Weigend, S.: Empirical evaluation of genetic clustering methods using multilocus genotypes from 20 chicken breeds. Genetics 159(2), 699–713 (2001)

53. Banks, M.A., Eichert, W., Olsen, J.B.: Which genetic loci have greater population assignment power? Bioinformatics 19(11), 1436–1438 (2003). doi:10.1093/bioinformatics/btg172

54. Rosenberg, N.A.: Algorithms for Selecting Informative Marker Panels for Population Assignment. JOURNAL OF COMPUTATIONAL BIOLOGY 12(9), 1183–1201 (2005)

55. Bromaghin, J.F.: Bels: Backward elimination locus selection for studies of mixture composition or individual assignment. Molecular Ecology Resources 8(3), 568–571 (2008). doi:10.1111/j.1471-8286.2007.02010.x

56. Slatkin, M.: Linkage disequilibrium — understanding the evolutionary past and mapping the medical future. Nature Reviews Genetics 9(6), 477–485 (2008). doi:10.1038/nrg2361

57. Cover, T.M., Thomas, J.A.: Elements of Information Theory. Wiley-Interscience, Hoboken, NJ, USA (2006)

58. EFSA: The European Union summary report on trends and sources ofzoonoses, zoonotic agents and food-borne outbreaks in 2015. EFSA Journal 14(12), 4634 (2016). doi:10.2903/j.efsa.2016.4634

59. Taylor, E.V., Herman, K.M., Ailes, E.C., Fitzgerald, C., Yoder, J.S., Mahon, B.E., Tauxe, R.V.: Common source outbreaks of Campylobacter infection in the USA, 1997-2008. Epidemiology and Infection 141(5), 987–996 (2013). doi: 10.1017/S0950268812001744

60. Li, J.Z., Absher, D.M., Tang, H., Southwick, A.M., Casto, A.M., Ramachandran, S., Cann, H.M., Barsh, G.S., Feldman, M., Cavalli-sforza, L.L., Myers, R.M.: Worlwide Human Relationships Inferred from Genome-Wide Patterns of Variation. Science 319(February), 1100–1104 (2008)

61. Huang, L., Jakobsson, M., Pemberton, T.J., Ibrahim, M., Nyambo, T., Omar, S., Pritchard, J.K., Tishkoff, S.A., Rosenberg, N.A.: Haplotype variation and genotype imputation in African populations. Genetic Epidemiology 35(8), 766–780 (2011). doi:10.1002/gepi.20626

62. Xuereb, A., Benestan, L., Normandeau, É., Daigle, R.M., Curtis, J.M.R., Bernatchez, L., Fortin, M.-J.: Asymmetric oceanographic processes mediate connectivity and population genetic structure, as revealed by RADseq, in a highly dispersive marine invertebrate (<i>Parastichopus californicus</i>). Molecular Ecology 27(10), 2347–2364 (2018). doi:10.1111/mec.14589

63. Tyanova, S., Albrechtsen, R., Kronqvist, P., Cox, J., Mann, M., Geiger, T.: Proteomic maps of breast cancer subtypes. Nature Communications 7, 1–11 (2016). doi:10.1038/ncomms10259

64. Lesk, A.M.: Introduction to Bioinformatics, 4th edn. Oxford University Press, Oxford (2014)

65. Efron, B.: Estimating the Error Rate of a Prediction Rule: Improvement on Cross-Validation. Journal of the American Statistical Association 78(382), 316 (1983). doi: 10.2307/2288636

66. Kuhn, M., Johnson, K.: Applied Predictive Modeling. Springer, New York, NY (2013). doi:10.1007/978-1-4614-6849-3. http://link.springer.com/10.1007/978-1-4614-6849-3

67. R Core Team: R: A Language and Environment for Statistical Computing, Vienna, Austria (2015). https://www.r-project.org/

68. Bansal, V., Libiger, O.: Fast individual ancestry inference from DNA sequence data leveraging allele frequencies for multiple populations. BMC Bioinformatics 16(1), 4 (2015). doi: 10.1186/s12859-014-0418-7

69. Nei, M.: Analysis of Gene Diversity in Subdivided Populations. Proceedings of the National Academy of Sciences 70(12), 3321–3323 (1973). doi: 10.1073/pnas.70.12.3321

70. Rannala, B., Mountain, J.L.: Detecting immigration by using multilocus genotypes. Proceedings of the National Academy of Sciences of the United States of America 94(17), 9197–201 (1997)

71. Wilson A., G., Rannala, B.: Bayesian inference of recent migration rates using multilocus genotypes. Genetics 163(3), 1177–1191 (2003). doi:Article./ehis.ebscohost.com/

72. >Mughini-Gras, L., Enserink, R., Friesema, I., Heck, M., van Duynhoven, Y., van Pelt, W.: Risk Factors for Human Salmonellosis Originating from Pigs, Cattle, Broiler Chickens and Egg Laying Hens: A Combined Case-Control and Source Attribution Analysis. PLoS ONE 9(2), 87933 (2014). doi:10.1371/journal.pone.0087933

73. Hald, T., Vose, D., Wegener, H.C., Koupeev, T.: A Bayesian Approach to Quantify the Contribution of Animal-Food Sources to Human Salmonellosis. Risk analysis 24(1), 255–269 (2004). doi:10.1111/j.0272-4332.2004.00427.x

74. Piry, S., Alapetite, A., Cornuet, J.-M., Paetkau, D., Baudouin, L., Estoup, A.: GENECLASS2: A Software for Genetic Assignment and First-Generation Migrant Detection. Journal of Heredity 95(6), 536–539 (2004). doi: 10.1093/jhered/esh074

75. Mughini-Gras, L., van Pelt, W.: Salmonella source attribution based on microbial subtyping: Does including data on food consumption matter? International Journal of Food Microbiology 191, 109–115 (2014). doi:10.1016/J.IJFOODMICRO.2014.09.010

76. Paetkau, D., Slade, R., Burden, M., Estoup, A.: Genetic assignment methods for the direct, real-time estimation of migration rate: a simulation-based exploration of accuracy and power. Molecular Ecology 13(1), 55–65 (2004). doi:10.1046/j.1365-294X.2004.02008.x

77. Andrews, K.R., Adams, J.R., Cassirer, E.F., Plowright, R.K., Gardner, C., Dwire, M., Hohenlohe, P.A., Waits, L.P.: A bioinformatic pipeline for identifying informative SNP panels for parentage assignment from RADseq data. Molecular Ecology Resources 18(6), 1263–1281 (2018). doi: 10.1111/1755-0998.12910

78. Freeland, J.R., Kirk, H., Petersen, S.: Molecular Ecology, 2nd edn. Wiley-Blackwell, Chichester, UK (2011)

79. Laing, C., Buchanan, C., Taboada, E.N., Zhang, Y., Kropinski, A., Villegas, A., Thomas, J.E., Gannon, V.P.: Pan-genome sequence analysis using Panseq: an online tool for the rapid analysis of core and accessory genomic regions. BMC Bioinformatics 11(1), 461 (2010). doi: 10.1186/1471-2105-11-461

80. Davidson, A.C., Hinkley, D.V.: Bootstrap Methods And Their Application. Cambridge University Press, Cambridge (1997)

81. Spiegelhalter, D.: The Art of Statistics: Learning from Data. Pelican, London (2019)

82. Wegner, P., Peter: A technique for counting ones in a binary computer. Communications of the ACM 3(5), 322 (1960). doi:10.1145/367236.367286

83. McCarthy, N.D., Colles, F.M., Dingle, K.E., Bagnall, M.C., Manning, G., Maiden, M.C.J., Falush, D.: Host-associated genetic import in Campylobacter jejuni. Emerging Infectious Diseases 13(2), 267–272 (2007). doi:10.3201/eid1302.060620

84. Lande, R.: Statistics and Partitioning of Species Diversity, and Similarity among Multiple Communities. Oikos 76(1), 5 (1996). doi: 10.2307/3545743. arXiv:1011.1669v3

85. Jost, L.: Entropy and diversity. Oikos 113(2), 363–375 (2006). doi:10.1111/j.2006.0030-1299.14714.x

86. Sherwin, W.B., B., W.: Entropy and Information Approaches to Genetic Diversity and its Expression: Genomic Geography. Entropy 12(7), 1765–1798 (2010). doi:10.3390/e12071765

87. Smith, R.D.: Information Theory and Population Genetics. arXiv:1103.5625 (2011). 1103.5625

88. Hu, H., Liu, X., Jin, W., Hilger Ropers, H., Wienker, T.F.: Evaluating information content of SNPs for sample-tagging in re-sequencing projects. Scientific Reports 5, 10247 (2015). doi:10.1038/srep10247

89. Liu, Z., Lin, S.: Multilocus LD measure and tagging SNP selection with generalized mutual information. Genetic Epidemiology 29(4), 353–364 (2005). doi: 10.1002/gepi.20092

90. Zhang, L., Liu, J., Deng, H.-W.: A multilocus linkage disequilibrium measure based on mutual information theory and its applications. Genetica 137(3), 355–364 (2009). doi:10.1007/s10709-009-9399-2

## References

1. Pemberton TJ, DeGiorgio M, Rosenberg NA. 2013 Population Structure in a Comprehensive Genomic Data Set on Human Microsatellite Variation. G3 Genesi/Genomes/Genetics 3, 891–907. (doi:10.1534/g3.113.005728)

2. Huang L, Jakobsson M, Pemberton TJ, Ibrahim M, Nyambo T, Omar S, Pritchard JK, Tishkoff SA, Rosenberg NA. 2011 Haplotype variation and genotype imputation in African populations. Genet. Epidemiol. 35, 766–780. (doi:10.1002/gepi.20626)

3. Li JZ et al. 2008 Worlwide Human Relationships Inferred from Genome-Wide Patterns of Variation. Science (80-.). 319, 1100–1104.

4. Xuereb A, Benestan L, Normandeau É, Daigle RM, Curtis JMR, Bernatchez L, Fortin M-J. 2018 Asymmetric oceanographic processes mediate connectivity and population genetic structure, as revealed by RADseq, in a highly dispersive marine invertebrate (*Parastichopus californicus*). Mol. Ecol. 27, 2347–2364. (doi:10.1111/mec.14589)

5. Tyanova S, Albrechtsen R, Kronqvist P, Cox J, Mann M, Geiger T. 2016 Proteomic maps of breast cancer subtypes. Nat. Commun. 7, 1–11. (doi:10.1038/ncomms10259)

## References

[1] A. Xuereb, L. Benestan, É. Normandeau, R. M. Daigle, J. M. R. Curtis, L. Bernatchez, and M.-J. Fortin, Molecular Ecology 27, 2347 (2018).

[2] M. Kuhn and K. Johnson, Applied Predictive Modeling (Springer New York, New York, NY, 2013).

## References

[1] S. Tyanova, R. Albrechtsen, P. Kronqvist, J. Cox, M. Mann, and T. Geiger, Nature Communications 7, 1 (2016).

[2] M. Kuhn and K. Johnson, Applied Predictive Modeling (Springer New York, New York, NY, 2013).

## References

[1] Y. H. Wang, Statistica Sinica 3, 295 (1993).

[2] B. Rannala and J. L. Mountain, Proceedings of the National Academy of Sciences 94 (1997).

[3] D. Paetkau, W. Calvert, I. Stirling, and C. Strobeck, Molecular Ecology 4, 347 (1995).

[4] J. M. Cornuet, S. Piry, G. Luikart, A. Estoup, and M. Solignac, Genetics 153, 1989 (1999).

[5] S. Piry, A. Alapetite, J.-M. Cornuet, D. Paetkau, L. Baudouin, and A. Estoup, Journal of Heredity 95, 536 (2004).

[6] L. Mughini-Gras and W. van Pelt, International Journal of Food Microbiology 191, 109 (2014).

[7] J. K. Pritchard, M. M. Stephens, and P. Donnelly, Genetics 155, 945 (2000).

[8] N. Takezaki and M. Nei, Genetics 144, 389 (1996).

[9] M. Nei, Proceedings of the National Academy of Sciences 70, 3321 (1973)

## References

[1] S. Manel, P. Berthier, and G. Luikart, Conservation Biology 16, 650 (2002).

